# Bidirectional modulation of negative emotional states by parallel genetically-distinct basolateral amygdala pathways to ventral striatum subregions

**DOI:** 10.1101/2024.06.19.599749

**Authors:** Sarah E. Sniffen, Sang Eun Ryu, Milayna M. Kokoska, Janardhan Bhattarai, Yingqi Wang, Ellyse R. Thomas, Graylin M. Skates, Natalie L. Johnson, Andy A. Chavez, Sophia R. Iaconis, Emma Janke, Minghong Ma, Daniel W. Wesson

## Abstract

Distinct basolateral amygdala (BLA) cell populations influence emotional responses in manners thought important for anxiety and anxiety disorders. The BLA contains numerous cell types which can broadcast information into structures that may elicit changes in emotional states and behaviors. BLA excitatory neurons can be divided into two main classes, one of which expresses *Ppp1r1b* (encoding protein phosphatase 1 regulatory inhibitor subunit 1B) which is downstream of the genes encoding the D1 and D2 dopamine receptors (*drd1* and *drd2* respectively). The role of *drd1+* or *drd2+* BLA neurons in learned and unlearned emotional responses is unknown. Here, we identified that the *drd1*+ and *drd2*+ BLA neuron populations form two parallel pathways for communication with the ventral striatum. These neurons arise from the basal nucleus of the BLA, innervate the entire space of the ventral striatum, and are capable of exciting ventral striatum neurons. Further, through three separate behavioral assays, we found that the *drd1*+ and *drd2*+ parallel pathways bidirectionally influence both learned and unlearned emotional states when they are activated or suppressed, and do so depending upon where they synapse in the ventral striatum – with unique contributions of *drd1*+ and *drd2*+ circuitry on negative emotional states. Overall, these results contribute to a model whereby parallel, genetically-distinct BLA to ventral striatum circuits inform emotional states in a projection-specific manner.

## Introduction

The ability to evaluate a sensory stimulus and to correctly act upon it is paramount for survival. Abnormal associations of stimuli and/or abnormal actions towards stimuli are hallmark features of psychiatric disorders including anxiety disorders. The basolateral amygdala (BLA) has long been known to support emotional responses (Klüver and Bucy, 1939; Blanchard and Blanchard, 1972), including to both aversive (Weiskrantz, 1956; Cahill and McGaugh, 1990; LeDoux, 1992; Maren et al., 1996; Cousens and Otto, 1998) and appetitive stimuli (Hatfield et al., 1996; Setlow et al., 2002; Schoenbaum et al., 2003). Affording the BLA with this capacity are both its intrinsic plasticity (Rogan et al., 1997; Maren and Quirk, 2004) and its projections into ‘downstream’ structures which can directly influence decisions and behavioral outcomes. For instance, the BLA innervates the central nucleus of the amygdala and this input is necessary for the expression of learned avoidance (LeDoux, 2003). Photostimulating central amygdala projecting BLA neurons evokes avoidance, while photoinhibition of the same neurons reduces fear learning (Namburi et al., 2015). Other projections of the BLA can influence appetitive responses, including stimulation of BLA neurons that project to the ventral striatum’s nucleus accumbens (NAc) (Namburi et al., 2015). Thus, it is clear that regionally-separable downstream recipients of BLA input are sufficient to direct emotional responses.

There is also an interplay between BLA projection targets and the cell types which comprise those projections in how specific BLA outputs influence emotion. The genetic identity of BLA neurons is highly diverse and these different cell types appear to be uniquely be engaged following emotional responses (Hochgerner et al., 2023). A single genetically distinct neuronal population can drive opposing emotional responses if it were to project to two brain regions (Zhang et al., 2021) and likewise, two genetically distinct BLA outputs can drive opposing emotional responses if they each project to the same brain region (Kim et al., 2017). BLA excitatory neurons are divided into two main genetic classes, which are becoming increasingly understood to have diverse projection targets and functions. These include the *Rspo2* expressing neurons (encoding R-spondin 2), which can drive aversive behaviors, and the *Ppp1r1b* expressing neurons (encoding protein phosphatase 1 regulatory inhibitor subunit 1B), which appear to support appetitive behaviors (Kim et al., 2016). BLA neurons distinguished by the expression of the transcription factor *Rspo2*, are also labeled by *Fezf2* (encoding the transcription factor zinc-finger 2) (Zhang et al., 2021), and project to the NAc and also its neighboring ventral striatum subregion, the tubular striatum (TuS, also known as the olfactory tubercle) (Wesson, 2020). Activation of *Fezf2* neurons innervating the NAc drives aversive states and contrastingly, activation of *Fezf2* neurons innervating the TuS increases appetitive states. Together, these findings indicate that neither the downstream target nor the genetic identity alone sufficiently explain how the BLA broadcasts emotional information. Instead, where this information goes *and* who within the BLA sends it are *both* critical for regulating emotional states. Given that *Rspo2*/*Fezf2*+ BLA neurons support both appetitive and aversive states depending upon their downstream targeting, we sought to answer the question of whether the *Ppp1r1b*+ BLA neuron population also contribute to aversive states, and do so depending upon their regional innervation within the NAc and TuS.

*Ppp1r1b* (also known as *Darpp-32,* dopamine- and cAMP-regulated neuronal phosphoprotein) is a phosphoprotein regulated by both D1 and D2 dopamine receptors (Ouimet et al., 1984; Nishi et al., 1997; Svenningsson et al., 2004), which are encoded for by the *drd1* and *drd2* genes, respectively (Scibilia et al., 1992). Dopamine within the BLA is necessary for fear learning (Fadok et al., 2009). We know that *drd1+* neurons in the BLA contribute to memory (Zhang et al., 2020a). While the role of *drd2*+ neurons in the BLA is not understood, prior pharmacological work has indicated a role for the D2 receptor in emotional responses (Guarraci et al., 2000; Berglind et al., 2006; de Oliveira et al., 2011). Overall, the respective contributions of *drd1+* and *drd2+* BLA neurons in regulating emotional states are unknown. Moreover, it is unknown if, like the *Fezf2* neurons, the influence of *drd1+* and *drd2+* BLA neurons depends upon their projection targets. Here, using a combination of viral tracing, *ex vivo* brain slice recordings, chemo- and opto-genetics, and behavior, we identified that the *drd1+* and *drd2+* BLA neuron populations form two parallel pathways wherein each innervate both the NAc and the TuS for the modulation of negative emotional states depending upon which ventral striatum subregion they innervate. Overall, these results contribute to a model whereby parallel, genetically-distinct, BLA to ventral striatum circuits inform emotional states in a projection-specific manner and altogether expand our appreciation for how the BLA regulates emotions.

## Results

### drd1+ and drd2+ BLA neurons innervate the ventral striatum

We first sought to determine if BLA *drd1+* and *drd2+* neurons form a circuit with ventral striatum neurons. We injected a Cre-dependent retrograde (rg) AAV expressing mCherry, rgAAV.hSyn.DIO.mCherry, into either the NAc or the TuS of *drd1-Cre* and *drd2-Cre* mice (Gong et al., 2007) (**Fig 1A,B & 1E,F**) and later inspected the BLA for mCherry+ neurons. mCherry+ cells in both groups of mice were found in the BLA in both *drd1-* and *drd2-Cre* mice (**Fig 1C & G**), indicating that these neurons indeed project to the ventral striatum. mCherry+ cells were found throughout the entire anterior-posterior extent of the BLA (**Fig 1C & G**). In contrast to the lateral amygdala (LA) which was largely void of mCherry+ cells, the basal nucleus of the amygdala (BA) displayed dense mCherry+ cells (**Fig 1C & G**). This organization was observed even when similarly injecting either the NAc or TuS of Ai9.TdTomato Cre reporter mice (Madisen et al., 2010) with rgAAV.hSyn.HI.eGFP-Cre.WPRE.SV40, suggesting the BA is a major conduit of ventral striatum input regardless of cell type (**Supplementary Fig 1A-C & 1D-F**). No reciprocal connection from ventral striatum to the BLA was found however (**Supplementary Fig 1G-I**).

**Figure 1.**
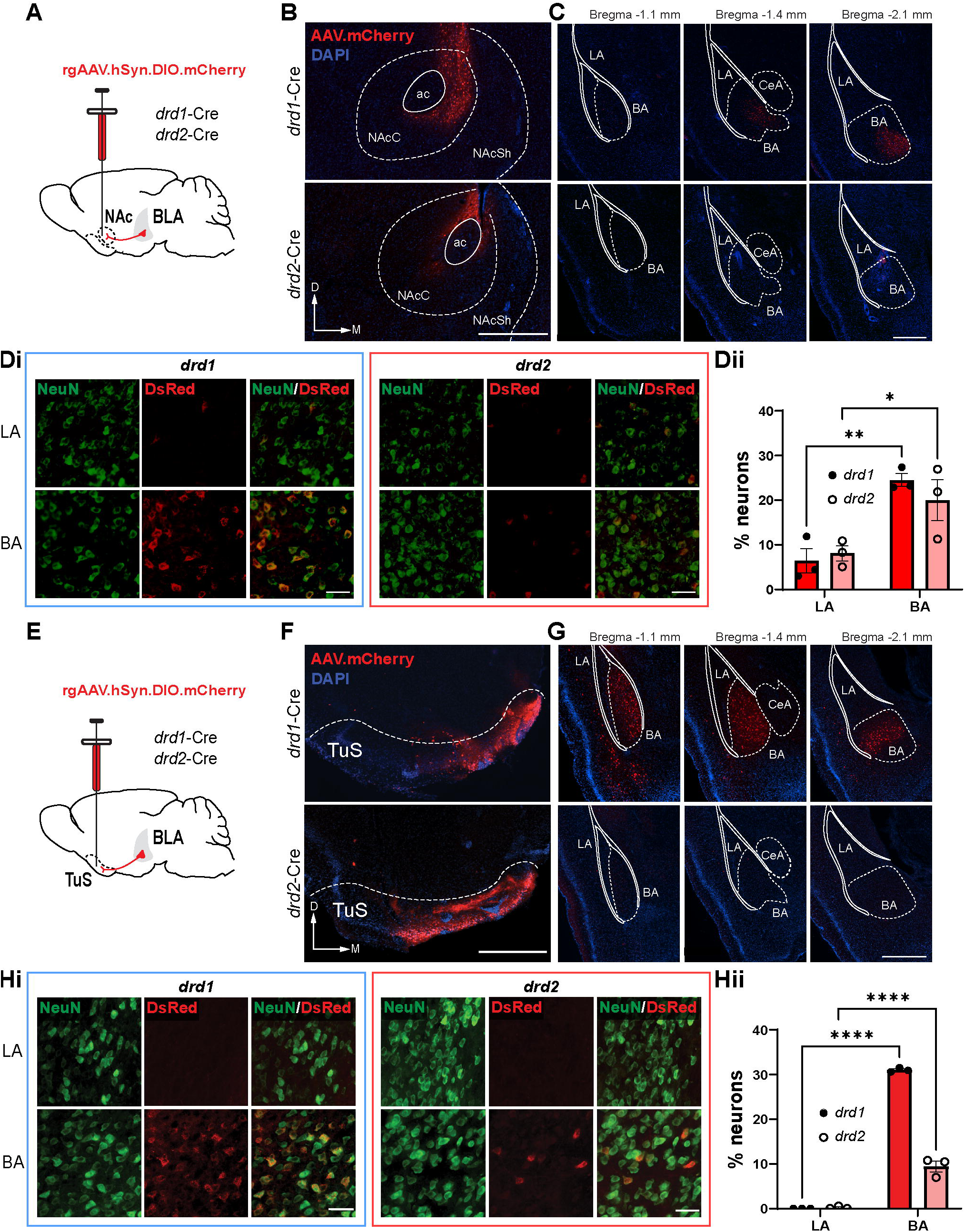
Ventral striatum projecting BLA neurons arise from the BA and are comprised of *drd1*+ and *drd2*+ neurons. **(A)** Schematic of approach for identifying BLA *drd1+* and *drd2+* inputs to the NAc. **(B)** Example of an NAc injection in *drd1*-Cre (top) and *drd2*-Cre (bottom) mice (ac = anterior commissure, NAcC & NAcSh=nucleus accumbens core & shell, respectively), and **(C)** example images of NAc projecting *drd1+* (top) or *drd2*+ (bottom) neurons along the anterior-posterior axis of the BLA. Scale bars=500µm. **(Di)** Anti-NeuN and anti-DsRed immunofluorescence images to identify the size of the NAc projecting BLA neural population. Scale bars=40µm. **(Dii)** Quantification of the *drd1* (n=3) or *drd2* (n=3) expressing BLA cells along the entire AP axis that project to the NAc (Two-way ANOVA, ROI main effect *F*(1,8)=26.65, *p*=0.001). **(E)** Schematic of approach for identifying BLA *drd1+* and *drd2+* inputs to the TuS. **(F)** Example of a TuS injection in *drd1*-Cre (top) and *drd2*-Cre (bottom) mice (TuS=tubular striatum), and **(G)** example images of TuS projecting *drd1+* (top) or *drd2*+ (bottom) neurons along the anterior-posterior axis of the BLA. Scale bars=500µm. **(Hi)** Anti-NeuN and anti-DsRed immunofluorescence images to identify the size of the NAc projecting BLA neural population. Scale bars=40µm. **(Hii)** Quantification of the *drd1* (n=3) or *drd2* (n=3) expressing TuS projecting BLA cells along the entire AP axis (Two-way ANOVA, *F*(1,8)=295.3, *p*<0.001). Mean±SEM.

To quantify the spatial distribution and to identify the size of the *drd1* and *drd2* neural population innervating both the NAc and TuS, sections were immunolabeled for both NeuN and the red fluorescent protein DsRed (**Fig 1D i & 1Hi**). This revealed that indeed the vast majority of ventral striatum projecting *drd1+* and *drd2+* neurons arise from the BA (**Fig 1D ii,** NAc: *drd1+* MLSD=|BA-LA|=18.07, *p*=0.002, *drd2+* MLSD=|BA-LA|=11.95, *p*=0.019; **Fig 1H ii,** TuS: *drd1+* MLSD=|BA-LA|=31.0, *p*<0.001, *drd2+* MLSD=|BA-LA|=9.24, *p*<0.001; MLSD=mean least square difference). Further, more *drd1+* BA neurons innervate the TuS than those that are *drd2+* (**Fig 1H ii,** TuS: MLSD=|*drd1*-*drd2*|=21.6, *p*<0.001).

Where throughout the ventral striatum do BLA neurons innervate? To answer this, we injected *drd1-Cre* and adora2a (*a2a)-Cre* mice into the BLA with an AAV encoding a synaptophysin.mRuby fusion protein (AAV.hSyn.FLEx.mGFP-2A- Synaptophysin-mRuby) (Herman et al., 2016) (Fig 2A). *A2a*-Cre mice were chosen for this and later anterograde AAV-based experiments to attempt to achieve optimal presynaptic expression in *drd2+* neurons. This resulted in mRuby+ puncta, indicative of BLA *drd1*+ or *a2a*+ neuronal synapses, throughout both the NAc and TuS (**Fig 2B**). As expected based upon the retrograde tracing results in **Figure 1**, *drd1* mRuby+ puncta were highly visible in comparison to *a2a*+ (**Fig 2B & C**). The mRuby+ puncta spanned the entire medial-lateral and anterior-posterior extents of the TuS, and were especially prominent in layer 2 (**Fig 2B**) which is the densest cell layer. mRuby+ puncta were also observed throughout the medial-lateral and anterior-posterior extents of the NAc, with comparable amounts in both the NAc core and shell (**Fig 2C**, *drd1+* MLSD=|NAcC- NAcSh|=0.035, p>0.999, *a2a+* MLSD=|NAcC-NAcSh|=0.241, p>0.999). It is notable, given its roles in associative learning including fear learning (Li et al., 2008; Wilson and Sullivan, 2011), that the ventral striatum receives more BLA *drd1*+ and less *a2a*+ neuronal input than the neighboring piriform cortex (**Fig 2C**, *drd1+*: *F*(1,16)=30.8, p<0.001; *a2a+*: *F*(1,14)=10.6, *p*=0.006). Together these tracing results establish that both *drd1+* and *drd2+* neurons, largely form the BA, innervate the entire span of the ventral striatum.

**Figure 2.**
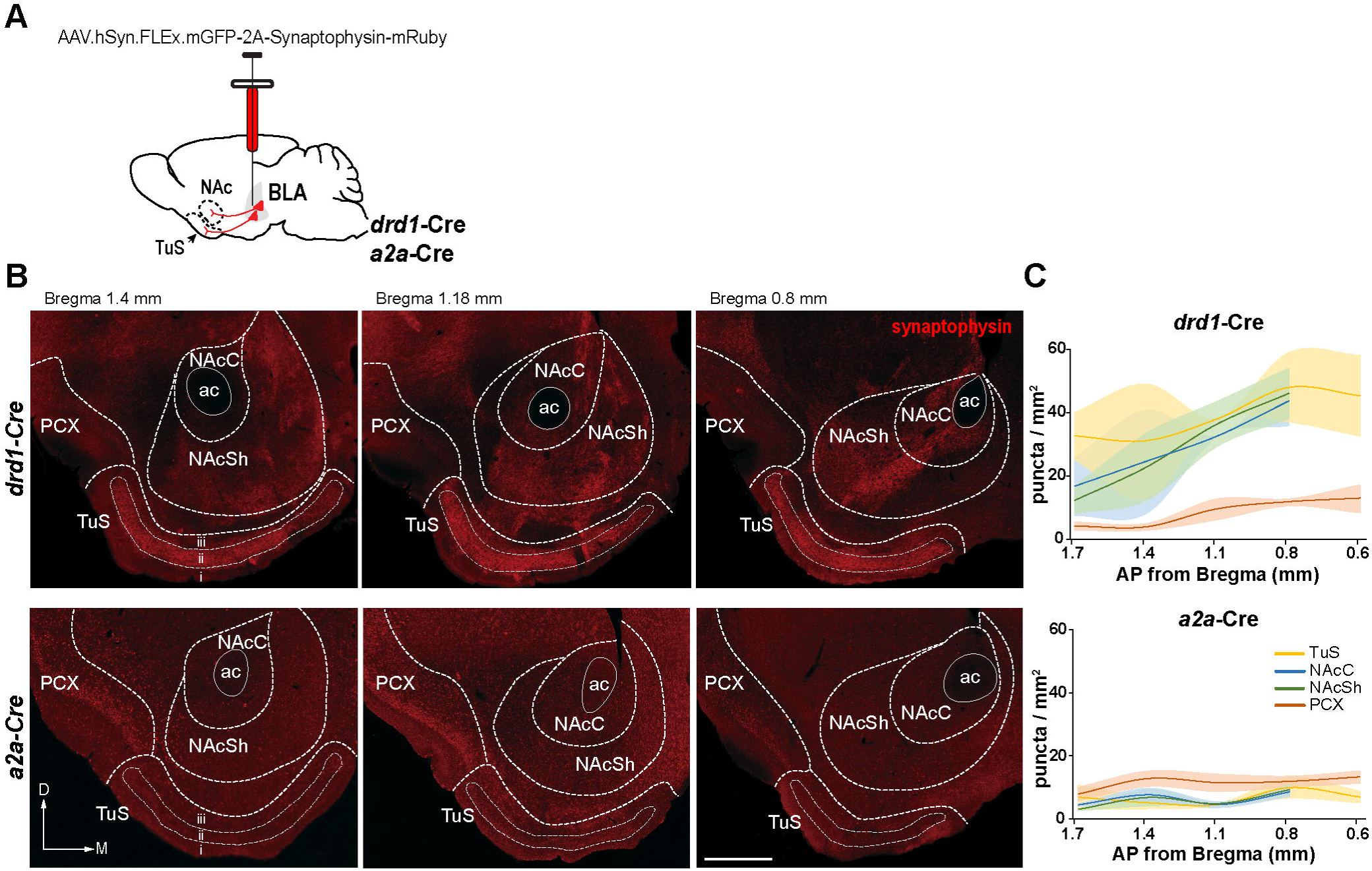
BLA neurons expressing *drd1* and *drd2* innervate the entire span of the ventral striatum. **(A)** Schematic of approach for identifying BLA *drd1+* and *drd2+* synaptic innervation of the ventral striatum. **(B)** Representative images showing direct innervation of BLA neurons into the ventral striatum in both *drd1*- and *a2a*-Cre mice. Scale bar=500µm. **(C)** Quantification of synaptophysin puncta in *drd1*-Cre (n=3) and *a2a*-Cre (n=3) mice. *drd1*+ BLA neurons densely innervate the ventral striatum, and relatively fewer *drd2*+ BLA neurons innervate the ventral striatum. Mean±SEM. PCX=piriform cortex, NAcC and NAcSh=nucleus accumbens core and shell, respectively.

### drd1+ and drd2+ BLA neurons excite ventral striatum spiny projection neurons

Next, we injected a Cre-dependent AAV expressing channelrhodopsin and EYFP (AAV.Ef1a.DIO.hChR2(E123T/T159C)-EYFP) or EYFP alone as a control (AAV.Ef1a.DIO.EYFP) into the BLA of *drd1-Cre* and *a2a-Cre* mice which were crossed with the Ai9 TdTomato Cre reporter line and later took coronal slices of the ventral striatum for *ex vivo* recordings to determine which ventral striatum spiny projection neurons (SPNs) the BLA neurons make synapses upon. TdTomato+ neurons were identified and used to identify *drd1+* or *drd2+* SPNs (those expressing TdTomato+ in either *drd1-Cre* or *a2a-Cre* mice respectively; **Fig 3A i & Aii**). In the same slices we also patched onto TdTomato- SPNs to monitor activity of *drd1*Ø and *drd2*Ø (putative *drd2+* or *drd1+*) SPNs, in either *drd1*-Cre or a2a-Cre mice respectively. During recordings, blue light pulses were delivered to excite ChR2-expressing BLA terminals in the ventral striatum. Importantly, we confirmed in *drd1-RFP;drd2-GFP* double transgenic mice (Shuen et al., 2008) that there is minimal co-expression of *drd1* and *drd2* in the same cells (**Supplementary Fig 2A & B).** Moreover, the BLA to ventral striatum projection is predominantly ipsilateral (**Supplementary Fig 2C & D**). Current injection into ventral striatum neurons confirmed their firing patterns are as expected for TuS SPNs (**Supplementary Fig 3A & B)** (White et al., 2019).

**Figure 3.**
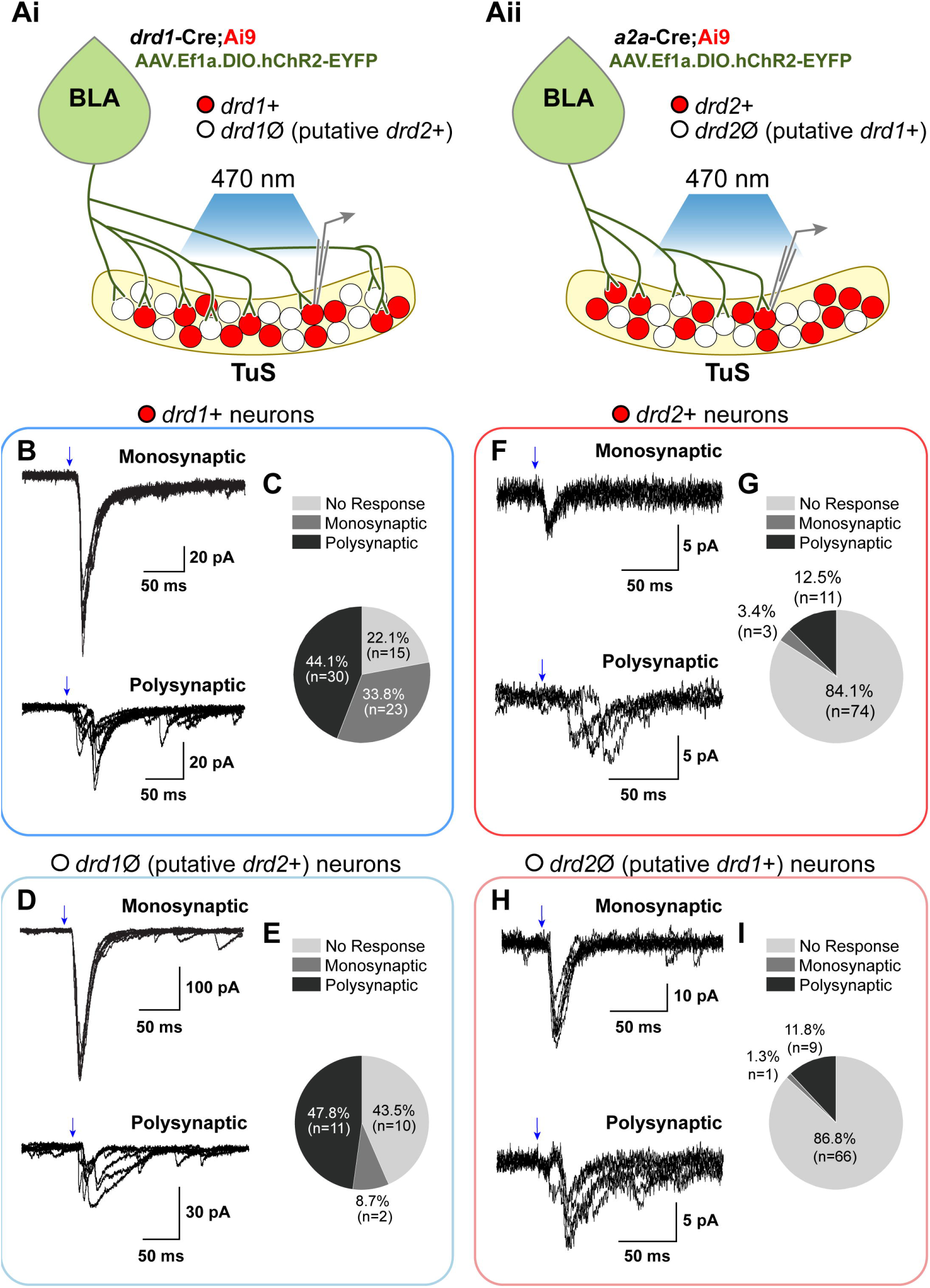
Synaptic properties of *drd1* and *drd2* expressing ventral striatum neurons receiving BLA neuronal projections. **(Ai)** Schematic indicating Cre-dependent expression of ChR2 in *drd1+* BLA neurons of *drd1*-Cre;Ai9 mice. In these mice, TdTomato+ neurons are presumably *drd1*+ and TdTomato- neurons are presumably *drd1*Ø. During whole-cell patch clamp recordings, ChR2 expressing BLA terminals were activated by 470nm light. **(Aii)** Schematic indicating Cre-dependent expression of ChR2 in *drd2+* BLA neurons of *A2a*-Cre;Ai9 mice. TdTomato+ neurons are presumably *drd2*+, and TdTomato- neurons are presumably *drd2*Ø. **(B)** Example light-evoked monosynaptic EPSCs (top) and light-evoked polysynaptic EPSCs (bottom) from *drd1*+ TuS neurons under voltage clamp mode. **(C)** Neurons organized by response type upon stimulation of *drd1*+ TuS projecting BLA terminals. **(D)** Example evoked EPSCs from *drd1*Ø TuS neurons. **(E)** Neurons organized by response type upon stimulation of *drd1*Ø TuS projecting BLA terminals. **(F)** Example evoked EPSCs from *drd2*+ TuS neurons. **(G)** Neurons organized by response type upon stimulation of *drd2*+ Tus projecting BLA terminals. **(H)** Example evoked EPSCs from *drd2*Ø TuS neurons. **(I)** Neurons organized by response type upon stimulation of *drd2*+ Tus projecting BLA terminals. The holding potential was −70mV.

We found that both BLA cell populations synapse upon both *drd1+* and *drd2+* SPNs, albeit with differing weights and strengths. The majority of *drd1+* BLA neurons elicited large monosynaptic currents in *drd1*+ and *drd1*Ø SPNs (**Fig 3B-D**). While both ventral striatum cell types were excited by *drd1*+ BLA neurons, BLA *drd1*+ neurons send stronger input (*viz.,* larger evoked excitatory postsynaptic currents [EPSCs]) and do so more predominantly upon *drd1+* vs *drd1*Ø (putative *drd2+*) SPNs (*drd1*+ vs *drd1*Ø monosynaptic: *X*^2^(1, N=91)=5.45, *p*=0.02; *drd1*+ vs *drd1*Ø polysynaptic: *X*^2^(1, N=91)=0.096, *p*=.757; **Fig 3C & E**). In a subset of SPNs, monosynaptic glutamatergic connections were verified via pharmacological manipulations (**Supplementary Fig 3C & D**). Likewise, *drd2+* BLA neurons also synapse upon *drd2+* and *drd2*Ø SPNs, but compared to the *drd1*+ BLA input, input from *drd2+* BLA neurons was both weaker and not as predominant (**Fig 3F-I**). A far larger percentage of SPNs displayed monosynaptic EPSCs upon *drd1*+ BLA neuron terminal stimulation than when stimulating *drd2*+ BLA terminals (*X*^2^(1, N=255)=33.947, *p*<0.001). Including polysynaptic EPSCs, 13.1 – 15.9% of SPNs (*drd2+* and *drd2*Ø, respectively) displayed evoked potentials and these were notably modest in amplitude compared to what was observed when stimulating *drd1*+ BLA terminals (e.g., **Fig 3B vs 3F**). Together these results extend the anatomical circuitry (**Figs 1 & 2**) by showing that both *drd1*+ and *drd2*+ BLA neurons excite ventral striatum spiny projection neurons, albeit with differing likelihood of observing monosynaptic connections.

### drd1+ BLA neurons innervating the NAc and drd2+ BLA neurons innervating the TuS both promote avoidance behavior

Next, we sought to determine a functional role for BLA *drd1+* and *drd2+* input to the ventral striatum. In the first of three assays, we used an optogenetic approach to excite *drd1+* and *drd2+* BLA neuron terminals innervating either the NAc or TuS to determine if these pathways influence avoidance or approach behaviors. For this we unilaterally injected *drd1*-Cre and *a2a*-Cre mice with AAV.Ef1a.DIO.hChR2(E123T/T159C)-EYFP or AAV.Ef1a.DIO.EYFP as control into their BLA and later implanted optical fibers into the ipsilateral NAc or TuS (**Fig 4A**). Given the similar innervation of both the NAc core and shell (**Fig 2C**), we targeted both NAc subregions for stimulation. Four weeks post injection, we used a 3-chamber real time place preference/aversion assay wherein light was delivered to stimulate the *drd1+* or *drd2+* BLA neuron terminals (465nm, 15ms pulses, 40Hz) on only one side of the chamber, with no optical stimulus in either the center or the opposite chambers (**Fig 4B**). The location of the mice was tracked with infrared photobeams to trigger the optogenetic stimulation and video was captured for off-line quantification.

**Figure 4.**
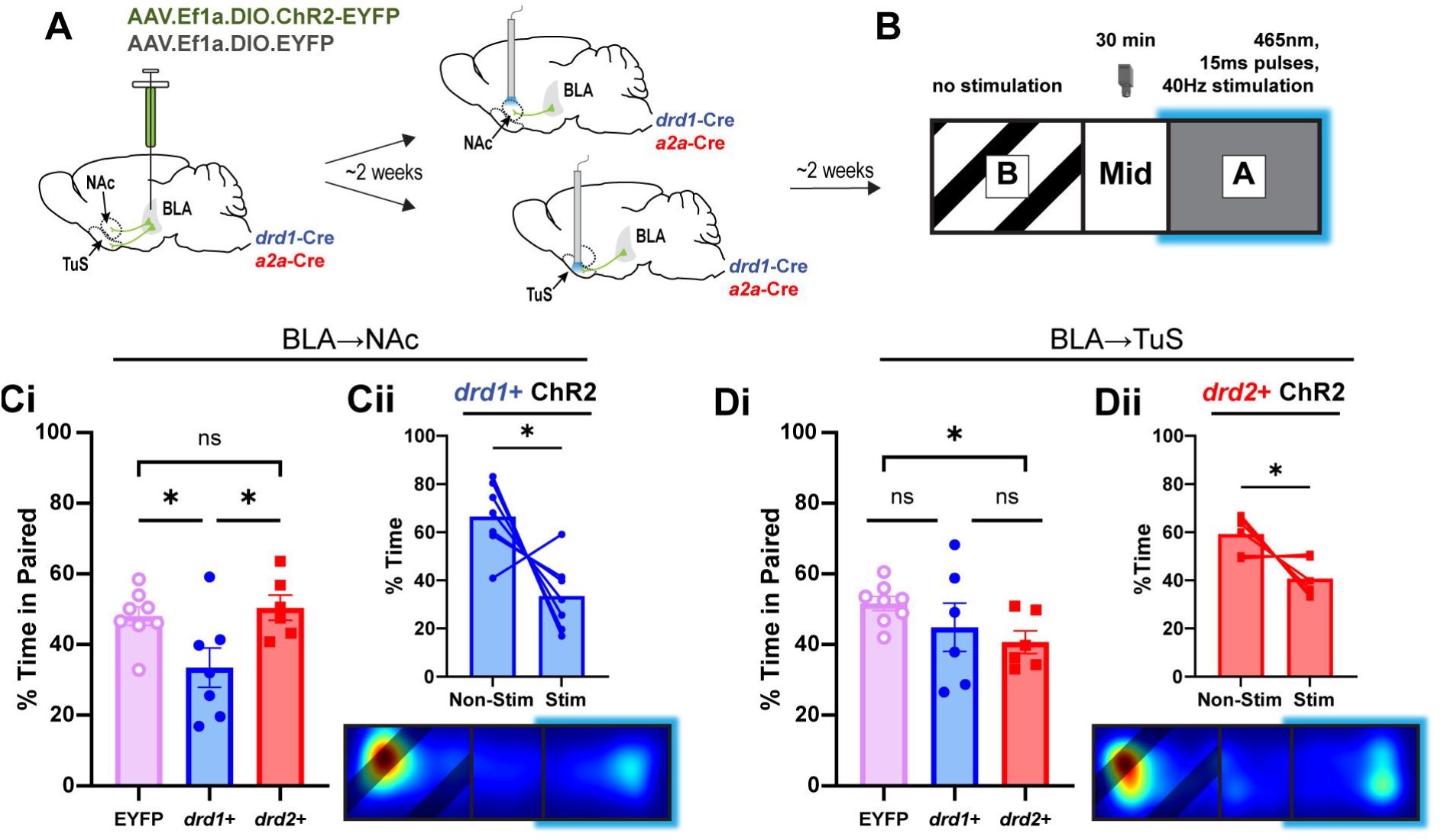
BLA *drd1+* and *drd2+* neurons innervating the ventral striatum promote aversive states depending upon projection target. **(A)** Paradigm for optic activation of NAc or TuS projecting *drd1+* or *drd2+* BA neurons and **(B)** 3-chamber real time place preference/aversion assay where optic stimulation occurs in only one side of the chamber (chamber A, blue glow). **(Ci)** Optical stimulation of *drd1+* BLA→NAc neurons resulted in less time spent in the light-paired chamber (One-way ANOVA, *F*(2,18)=5.04, *p*=0.018). **(Cii)** Stimulation of *drd1+* BLA→NAc neurons results in avoidance of the light- paired chamber (upper, *t*(6)=2.981, *p*=0.025), demonstrated by representative heat map of chamber preference from one mouse (lower). **(Di)** Optical stimulation of *drd2+* BLA→TuS neurons resulted in less time spent in the light-paired chamber compared to optical stimulation of EYFP controls (Welch’s ANOVA *W*(2.00,8.31)=6.02, *p*=0.024). **(Dii)** Stimulation of *drd2+* BLA→TuS neurons results in avoidance of the light-paired chamber (upper, *t*(5), *p*=2.916), demonstrated by representative heat map of chamber preference from one mouse (lower). Mean±SEM.

We found that optical stimulation of *drd1+* BLA➔NAc neuron terminals resulted in less time spent in the light-paired chamber compared to optical stimulation of *drd2+* BLA➔NAc neuron terminals and EYFP controls (**Fig 4C i & Cii**) (One-way ANOVA *F*(2,18)=5.04, *p*=0.018). Indeed, compared to the non-stimulated side, mice spent 49.70±11.10 % (mean ± SEM) less time on the chamber paired with *drd1+* BLA➔NAc neuron terminal stimulation (*t*(6)=2.981, *p*=0.025). Similarly, we found that optical stimulation of *drd2+* BLA➔TuS neuron terminals resulted in less time spent in the light-paired chamber compared to optical stimulation of *EYFP+* BLA➔TuS controls (**Fig 4D i & Dii**) (Welch’s ANOVA *W*(2.00,8.31)=6.02, *p*=0.024). Compared to the non-stimulated side, mice spent 31.42±6.39 % (mean ± SEM) less time on the chamber paired with *drd2+* BLA➔TuS neuron terminal stimulation (*t*(5)=2.916, *p*=0.033). These results show that activation of *drd1*+ and *drd2*+ BLA input to the NAc and TuS respectively lead to avoidance behavior.

### drd1+ BLA neurons innervating the NAc and drd2+ BLA neurons innervating the TuS support Pavlovian fear learning

Next, we wanted to know the possible influence of these pathways on learned emotional behaviors. To do this we employed an odor-shock Pavlovian fear-learning paradigm (Best and Wilson, 2003; Jones et al., 2008; Hegoburu et al., 2011) wherein an otherwise neutral odor is paired with a mild foot shock (**Supplementary Fig 4Ai**). To quantify learning, we monitored both physical immobility and fear-associated respiratory power (4-6Hz) which increases in power when animals anticipate an aversively-paired stimulus (Hegoburu et al., 2011; Moberly et al., 2018). Mice were placed in a plethysmograph with a custom floor made out of metal connected to a shock stimulus generator. Also connected to the plethysmograph was a tube allowing delivery of clean air or an odor which were both controlled by an odor presentation machine. All behavioral measures and stimuli were controlled by the same computer allowing synchrony in measures and stimulus presentation events. In untreated C57BL/6J mice we validated that only odors paired with shock, were associated with elevations in physical immobility following the conditioning day (**Supplementary Fig 4B**). We also validated that fear-associated 4-6Hz respiratory power is similarly elevated as mice learn to associate an odor with a shock (**Supplementary Fig 4C-H**).

We used a chemogenetic approach to suppress BLA➔ventral striatum input which included six separate groups of mice to establish the roles of each of the BLA➔NAc and BLA➔TuS pathways (**Fig 5A**). These included *drd1*+ and *drd2*+ mice injected with rgAAV.hSyn.DIO.hM4D(Gi)-mCherry or rgAAV.hSyn.DIO.mCherry as control. All mice were subsequently implanted with bilateral intracranial cannulae into the BLA for administration of either the DREADD ligand J60 (Bonaventura et al., 2019) or vehicle. J60 or vehicle were administered 30 minutes prior to the learning session following a single behavioral session on a prior day to acclimate the mice to the chamber.

**Figure 5.**
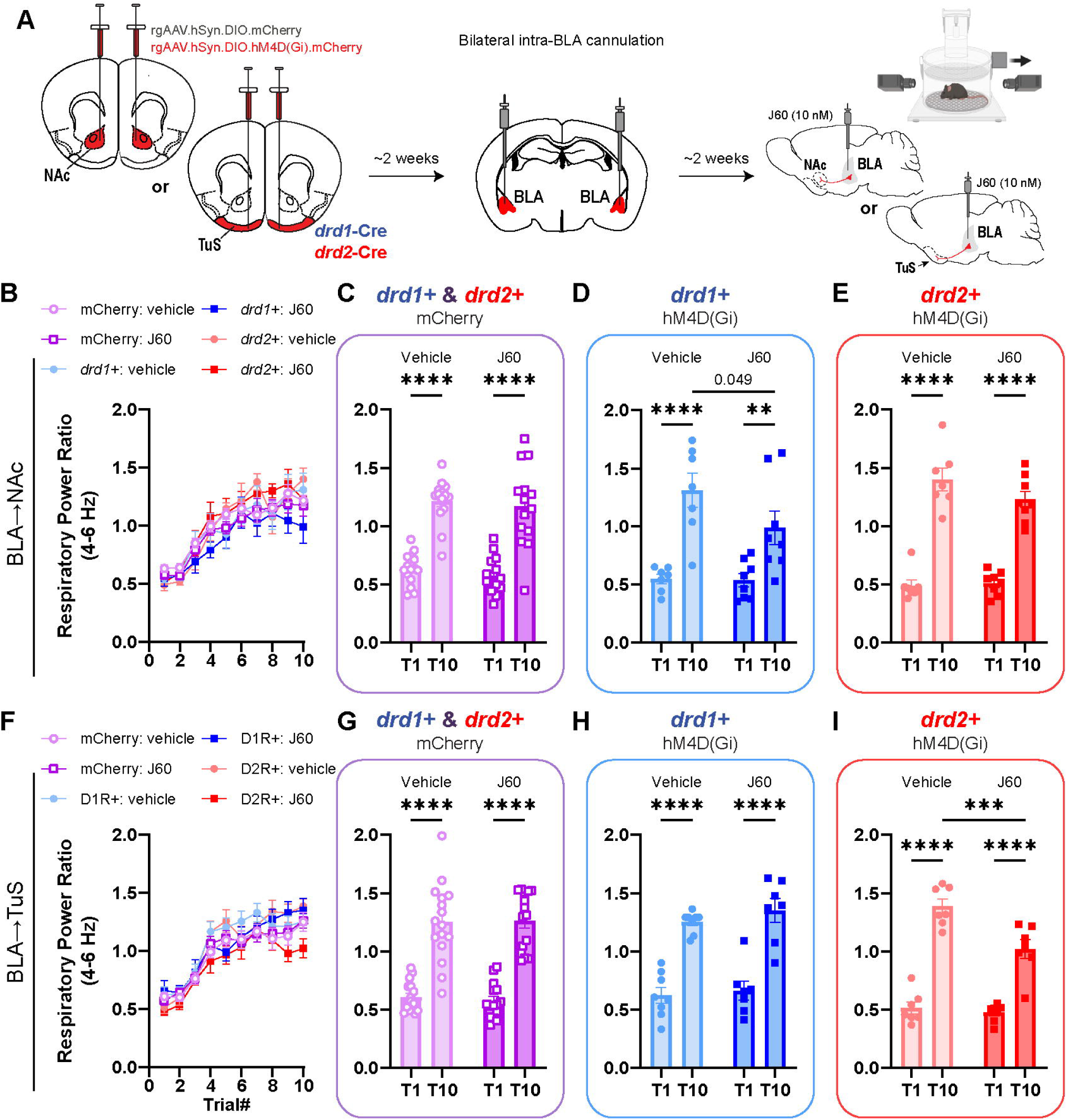
BLA *drd1+* and *drd2+* neurons innervate the ventral striatum support Pavlovian fear learning depending upon projection target. **(A)** Paradigm for DREADD induced silencing of NAc or TuS projecting *drd1*+ or *drd2*+ BLA cells. **(B)** Influence of DREADD agonist J60 (100nL, 10nM) on fear learning in all NAc injected mice, **(C)** left NAc injected mCherry controls (Two-way RM ANOVA, Trial main effect *F*(1,27)=86.2, *p*<0.001); middle *drd1*+ hM4D(Gi) mice (Two-way RM ANOVA, Trial main effect *F*(1,13)=48.9, *p*<0.001); and right *drd2*+ hM4D(Gi) mice (Two-way RM ANOVA, Trial main effect *F*(1,13)=301, *p*<0.001). **(D)** Influence of DREADD agonist J60 (100nL, 10nM) on fear learning in all TuS injected mice, **(E)** left mCherry TuS injected controls (Two-way RM ANOVA, Trial main effect *F*(1,29)=151, *p*<0.001); middle *drd1*+ hM4D(Gi) mice (Two-way RM ANOVA, Trial main effect *F*(1,13)=85.6, *p*<0.001); and right *drd2*+ hM4D(Gi) mice (Two-way RM ANOVA, *F*(1,12)=7.71, *p*=0.017). Mean±SEM.

Among both the BLA➔NAc and BLA➔TuS groups, all control groups displayed elevations in fear-associated respiration by the 10^th^ trial of odor-shock pairings (**Fig 5B,C & F,G**; NAc mCherry control: Two-way RM ANOVA, trial main effect *F*(1,27)=86.2, *p*<0.001; TuS mCherry control: Two-way RM ANOVA, trial main effect *F*(1,29)=151, *p*<0.001) indicating that they learned to associate an odor with an aversive outcome. As expected, similar elevations in physical immobility were also observed (**Supplementary Fig 4**). Importantly, there was no difference in learning between the vehicle and J60 infused groups supporting that there are no off-target effects of this DREADD ligand on odor-shock learning (**Fig 5B,C & F,G**; NAc mCherry controls Trial 10: MLSD=|Vehicle- J60|=-0.0443, *p*=0.601, TuS mCherry controls Trial 10: MLSD=|Vehicle-J60|=0.0118, *p*=0.888).

While neither inhibition of *drd2+* BLA➔NAc and *drd1+* BLA➔TuS pathways impacted fear-learning (**Fig 5E & H**, Two-way RM ANOVA, trial main effect: *drd2*+ BLA➔NAc: *F*(1,13)=301, *p*<0.001; *drd1*+ BLA➔TuS: *F*(1,13)=85.6, *p*<0.001), we found that inhibition of *drd1+* BLA➔NAc and *drd2+* BLA➔TuS pathways suppressed the magnitude of the learned association. Both *drd1+* BLA➔NAc and *drd2+* BLA➔TuS pathway inhibition resulted in less fear-related respiration by trial 10 in J60 infused mice compares to those infused with vehicle (**Fig 5D & I**; *drd1*+ BLA➔NAc Trial 10: MLSD=|Vehicle-J60|=0.319, *p*=0.049, *drd2*+ BLA➔TuS Trial 10: MLSD=|Vehicle- J60|=0.364, *p*<0.001). Fear-related physical immobility was likewise reduced upon *drd2+* BLA➔TuS pathway inhibition, yet interestingly not upon *drd1+* BLA➔NAc inhibition (**Supplementary Fig 5**). These results show that *drd1+* and *drd2+* BLA input to the NAc and TuS respectively are necessary for fear learning in addition to their role in real-time avoidance.

### drd2+ BLA neurons innervating the TuS promote spontaneous avoidance of odors

Finally, given the evidence that BLA to ventral striatum *drd1+* and *drd2+* neural pathways each influence both spontaneous / real-time aversive states and learned avoidance to odors, we examined if this circuitry might also influence spontaneous attraction or avoidance to odors. For this we used the same mice that completed the Pavlovian odor-shock fear learning (**Fig 5**), either a few days before or after the Pavlovian testing, and tested them in a spontaneous odor attraction/avoidance assay following intracranial infusion of either vehicle or J60. One side of the testing arena contained cotton laced with the aversive fox odor compound 2MT (2-Methyl-2- thiazoline), and the other side contained cotton laced with an attractive peanut oil odor (**Fig 6A**). Both stimuli were housed in clean perforated plastic tubes to prevent touching or tasting the stimulus yet still allowing release of volatiles. Video was collected for quantification of place preference. C57BL/6J mice, whether injected with J60 or vehicle, both spent more time on the peanut scented side of the apparatus than the fox odor side indicating that this approach can assay unlearned valence to odors (**Fig 6B**; vehicle: *t*(7)=3.71, *p*=0.004; J60: *t*(6)=2.24 *p*=0.033).

**Figure 6.**
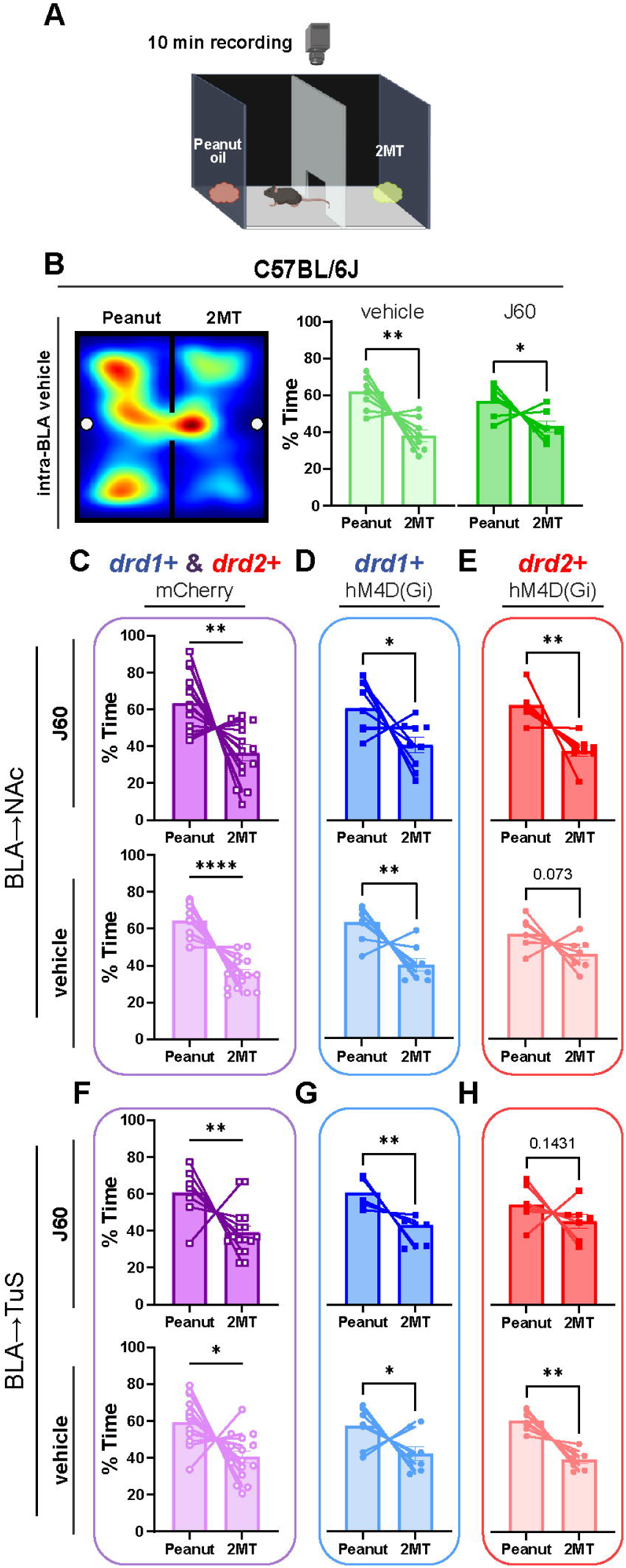
*drd2*+ BLA neurons innervating the TuS promote spontaneous avoidance to odors. **(A)** Behavioral schematic of spontaneous odor attraction/avoidance task. **(B-H)** Influence of DREADD agonist J60 (100nL, 10nM) on spontaneous avoidance in **(B)** C57BL/6J mice (vehicle: *t*(7)=3.71, *p*=0.004; J60: *t*(6)=2.24, *p*=0.033), **(C)** BLA→NAc mCherry control mice (top J60: *t*(13)=3.13, *p*=0.004; bottom vehicle: *t*(14)=6.31, *p*<0.001, **(D)** top J60 *drd1*+: *t*(8)=2.30, *p*=0.025; bottom vehicle *drd1*+: *t*(9)=3.840, *p*=0.002, **(E)** top J60 *drd2*+: *t*(7)=4.4, *p*=0.002; bottom right trend of *drd2*+ vehicle: *t*(6)=1.7, *p*=0.073). **(F)** BLA→TuS mCherry control mice (top J60: *t*(15)=3.34, *p*=0.002; bottom vehicle: *t*(14)=2.89, *p*=0.012, **(G)** top J60 *drd1*+: *t*(6)=3.705, *p*=0.005; bottom vehicle *drd1*+: *t*(7)=2.014, *p*=0.042, **(H)** top J60 *drd2*+: *t*(6)=1.171, *p*=0.143; bottom vehicle *drd2*+: *t*(6)=5.02, *p*=0.001). Mean±SEM.

As anticipated, among both the BLA➔NAc and BLA➔TuS mCherry groups, all controls, regardless of J60 or vehicle treatment, spent more time on the peanut scented side of the apparatus compared to the fox odor side (**Fig 6C & F**) supporting that there are no off-target effects of this DREADD ligand on spontaneous approach or avoidance to odors (J60-inhibited BLA➔NAc: *t*(13)=3.13, *p*=0.004; vehicle-treated BLA➔NAc: *t*(14)=6.31, *p*<0.001; J60-inhibited BLA➔TuS: *t*(15)=3.34, *p*=0.002; vehicle-treated BLA➔TuS: *t*(14)=2.89, *p*=0.012). In line with our results from the Pavlovian odor-shock fear learning, we found that *drd2+* BLA➔TuS pathway inhibition resulted in reduced approach and avoidance for peanut and fox, respectively (**Fig 6H**; J60-inhibited D2R+ BLA➔TuS: *t*(6)=1.171, *p*=0.143). Unlike in the Pavlovian odor-shock fear learning however, inhibition of *drd1+* BLA➔NAc pathway did not influence spontaneous approach and avoidance (**Fig 6D**) (J60-inhibited *drd1+* BLA➔NAc: *t*(8)=2.300, *p*=0.025). These results show that *drd2+* BLA input to the TuS, but not *drd1+* input to the NAc, contributes to unlearned odor avoidance in addition to its role in real-time avoidance and Pavlovian fear learning.

## Discussion

It is well established that BLA outputs to specific brain regions influence emotional responses (e.g., (Cardinal et al., 2002; Paré et al., 2004; Ambroggi et al., 2008; Stuber et al., 2011; Janak and Tye, 2015; Namburi et al., 2015; Beyeler et al., 2016)). More recently, several lines of evidence have uncovered divergent valence responding through genetically-distinct neurons within the BLA, including by means of *Pppr1r1b* and *Rspo2* neurons (Kim et al., 2016, 2017; Zhang et al., 2021). Together, both the genetic identity and downstream targets of BLA neurons are necessary to incorporate when understanding the role of BLA cell types in orchestrating the many functions of the BLA.

In the present study we focused on defining the contributions of *drd1*+ and *drd2*+ BLA neurons to emotional responding. The *drd1* and *drd2* genes encode for the D1 and D2 receptors, respectively (Scibilia et al., 1992), which regulate *Pppr1r1b –* a marker for one of the two main classes of BLA excitatory neurons. It has been long known that D1 and D2 receptors are in the BLA (e.g., (Meador-Woodruff et al., 1991)). We established that both *drd1*+ and *drd2*+ BLA neurons innervate the NAc and TuS, and we subsequently focused upon these two pathways (BLA→NAc and BLA→TuS) given the recent evidence of regulation of emotional responses through BLA output into these regions (Zhang et al., 2021). Our findings extend the work of (Zhang et al., 2021) by showing that in addition to the *Rspo2/Fezf2* BLA neuron class, both *drd1*+ and *drd2*+ BLA neurons in the *Ppp1r1b* neuron class also each innervate the NAc and TuS. Far more *drd1*+ neurons comprise this circuit than *drd2*+ neurons, with more *drd1+* neurons innervating both the TuS and NAc. These neurons originate from nearly the entire anterior-posterior extent of the BLA, and specifically the vast majority from within the BA (**Fig 1**). Further, our synaptophysin tracing suggests that they innervate nearly all of TuS and NAc space (all layers of TuS and both the NAc core and shell; **Fig 2**). While the spatial innervation of *drd1+* and *drd2+* BLA neurons into the ventral striatum is not unlike that reported by (Zhang et al., 2021), it is important to emphasize that *Fezf2*+ BLA neurons do not co-express *Pppr1r1b* (Zhang et al., 2021) which suggests these three neuron types connecting the BLA with the ventral striatum are distinct.

While the synaptophysin tracing suggests synaptic innervation of the ventral striatum by *drd1+* and *drd2+* BLA neurons, we used brain slice recordings to quantify this. This is interesting given that the primary cell type in the ventral striatum are spiny projection neurons which also express *drd1+* and *drd2+*. We focused our recordings on TuS spiny projection neurons given the comparable innervation of both structures (NAc and Tus) by *drd1+* and *drd2+* BLA neurons which allowed us to also perform recordings to identify if there is logic by which ventral striatum neurons these BLA neurons synapse upon. We found that both BLA cell types excite *drd1*+ and *drd2*+ (identified in this experiment by expression of A2a) TuS neurons in manners which appeared to be largely glutamatergic, with especially *drd1*+ BLA neurons sending a large amount of monosynaptic currents (**Supplementary Fig 3**). Further, *drd1+* BLA neurons monosynaptically excited predominately *drd1+* TuS neurons, and *drd2+* BLA neurons non-preferentially excited a small population of both *drd1+* and *drd2+* TuS neurons. While these results were initially surprising given that the *drd1*+ BLA→TuS pathway was dispensable for the fear and avoidance behaviors explored in this work, this may indicate a potential role for this pathway in other behaviors, such as those involved in reward. Thus, BLA input to the TuS, and therefore possibly also the NAc, has an organization which allows for recruitment of specific postsynaptic neurons in the TuS which could therefore allow differential output from the ventral striatum in manners supporting specific outputs into the basal ganglia and other brain networks important for behavioral responses.

We found within this circuitry that the parallel pathways generated by the *drd1+* and *drd2+* BLA neuron populations modulates negative emotional states depending upon their ventral striatum projection target (**Fig 7**). To show this, we used three distinct behavioral paradigms in combination with either projection specific chemo- or optogenetic manipulations. In all behavioral paradigms, we were able to uncover a role for either *drd1*+ and/or *drd2*+ neurons, yet, in not all cases did each cell population impact behavior. Instead, the impact on behavior was in most cases also projection target specific. For instance, *drd1*+ BLA neurons innervating the NAc increased negative valence states in the real-time place preference/aversion paradigm, whereas the same cell population projecting to the TuS did not (**Fig 4**). Likewise, *drd2*+ BLA neurons innervating the TuS increased negative valence states in the real-time place preference/aversion paradigm, whereas the same cell population projecting to the NAc did not. Similar differences in how these genetically-distinct BLA cell populations influenced Pavlovian fear learning and spontaneous valence behaviors were also observed to be cell-type and projection target specific. As mentioned, not in all cases did each pathway impact one of the three behaviors assayed, possibly hinting towards a role for these pathways in other behaviors. These findings lay the foundation for future work to systematically target *drd1*+ or *drd2*+ BLA inputs into specific regions within the NAc (core vs shell (West and Carelli, 2016)) or TuS (medial vs lateral (Murata et al., 2015; Zhang et al., 2017)) which may provide even more specific behavioral outcomes. This work extends a role for *drd1+* BLA neuron output to the central amygdala which was found to influence extinction memory (Zhang et al., 2020b), into two ventral striatum subregions which are important for valence-based behavioral responses, and by allowing for comparison with the influence of the neighboring *drd2+* neurons.

**Figure 7.**
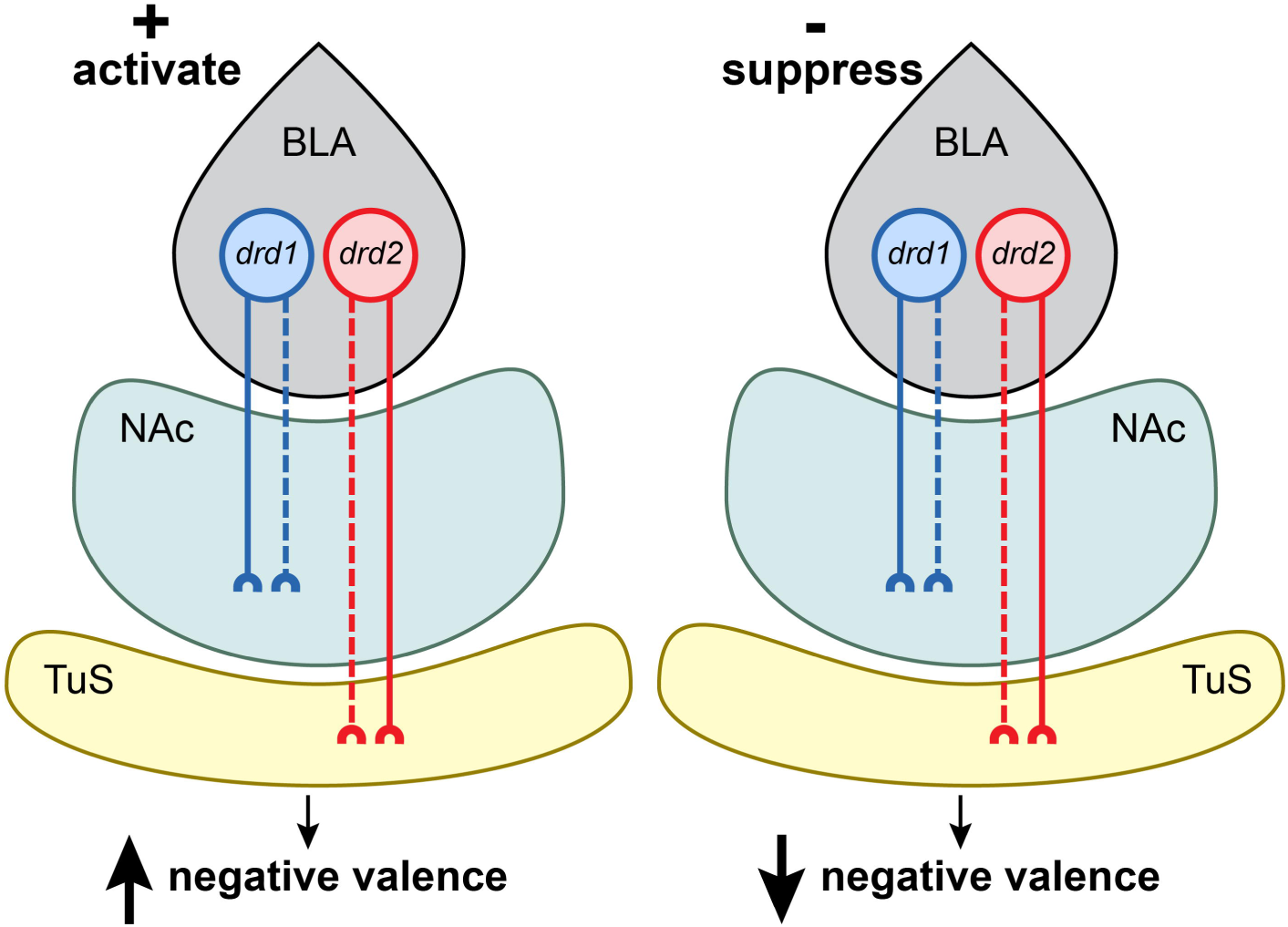
Overview of findings illustrating the behavioral consequences of activating (left) or suppressing (right) *drd1*+ or *drd2*+ BLA neuron inputs to the NAc and TuS. Dotted lines annotate pathways that are dispensable for regulating the behaviors tested.

Interestingly, when comparing changes in fear-associated respiration (well known to be influenced by sympathetic state (Stevenson and Ripley, 1952; Boiten, 1998; Homma and Masaoka, 2008; Hegoburu et al., 2011; Moberly et al., 2018)) and fear-induced immobility, we saw that manipulation of NAc projecting *drd1*+ BLA neurons did not similarly influence both of those fear-associated behaviors (**Supplementary Fig 5**). This may be due to either distinct inputs or outputs (collaterals) of the *drd1*+ BLA neurons which might differentially guide changes in respiratory behavior versus motor behavior. For instance, differential innervation of the periaqueductal grey might allow for one cell-type to influence respiration over another given the periaqueductal grey’s influence on breathing (Walker and Carrive, 2003; Subramanian and Holstege, 2013). While we did not identify the differential pathway, it is interesting to identify instances wherein fear-related behaviors are not simultaneously displayed. Further manipulation of this pathway while similarly taking multiple measures of fear-related behaviors, including even heart rate, skin conductance, and ultrasonic vocalizations for instance, will help refine our understanding of circuitry which specifically supports each to be displayed during emotional contexts.

Top-down glutamatergic inputs to the BLA profoundly influence associative learning and behaviors. Given the fact that these BLA→ventral striatum neurons express dopamine receptors, it is tempting to speculate how this pathway may be modulated by dopamine. Dopamine within the BLA is necessary for fear learning (Fadok et al., 2009). Local antagonism of both D1Rs and D2Rs within the BLA blocks the expression of fear during a potentiated startle paradigm (Lamont and Kokkinidis, 1998; Greba et al., 2001). Antagonism of BLA D1Rs also perturbs the timing of fear behavior (Shionoya et al., 2013), and antagonism of BLA D2Rs attenuates freezing during Pavlovian fear conditioning (Guarraci et al., 2000; de Oliveira et al., 2011; de Souza Caetano et al., 2013). The role of dopamine receptors is similarly mixed in appetitive behaviors, where antagonizing both D1Rs and D2Rs within the BLA attenuates conditioned reward seeking and taking (See et al., 2001; Berglind et al., 2006; Kim and Lattal, 2019). Indeed, local application of D1 agonists increases intrinsic excitability and the evoked firing of BLA neurons (Kröner et al., 2005). D1 receptors have a lower affinity for dopamine than D2 receptors (Richfield et al., 1989; Schultz, 2007). Further, when dopamine levels are low, D2 receptors are agonized, but when DA levels are elevated, like when receiving an emotionally salient stimulus, both D1 and D2 receptors become agonized (Guarraci et al., 1999; Horvitz, 2000; Bristol et al., 2004). It is possible these differential roles of D1 and D2 receptors in the BLA might explain our finding that *drd2*+ neurons vs *drd1*+ neurons contributed differently to the regulation of emotional states.

Overall, this work has uncovered that *drd1+* and *drd2+* neurons within the *Ppp1r1b* BLA neuron class forms parallel pathways which bidirectionally influence emotional states when they are activated or suppressed and do so depending upon where they synapse – with unique contributions of *drd1+* and *drd2+* BA→NAc vs BA→TuS circuitry on negative valence states. Overall, our results contribute to a model whereby parallel, genetically-distinct BLA to ventral striatum circuits inform emotional states in a projection-specific manner. This work adds to our understanding of the complex interplay between projection cell types and their projection targets, in how the BLA helps orchestrate emotions.

## Conflict of interest statement

The authors declare no competing financial interests.

## Acknowledgements

We thank Dr. Marc Fuccillo for generously sharing reagents for synaptophysin-based AAV tracing. This work was supported by R01DC014443 to D.W., and R01DA049545, and R01DC016519 to D.W.W and M.M.. N.L.J. was supported by NIDCD T32015994 and F31DC020364.. S.E.S. was supported by NIDCD T32015994 and F31DC02188801.

## Data analysis

Data were analyzed for statistical significance in GraphPad Prism. All data are reported as mean±SEM unless otherwise noted. Specific tests used can be found in the Results sections or the figure legends. All *t*-tests were paired. When possible, experimenters handling the data were blinded to treatment conditions.

## Materials and Methods

### Animals

Adult male and female mice, 2-5 months of age, were housed in a temperature-controlled vivarium on a 12:12 hour (hr) light/dark cycle with *ad libitum* access to food and water, except during behavioral testing. All behavioral testing occurred during the light cycle. Mice that only underwent viral injections were group housed (≤5 mice/cage) and mice with chronic implants were single housed following surgery. All experimental procedures were conducted within the AALAC animal research program of the University of Florida in accordance with the guidelines from the National Institute of Health, and were approved by the University of Florida Institutional Animal Care and Use Committee.

Mouse lines included the following transgenic lines which were maintained on a C57BL/6J background (strain #000664; RRID:IMSR_JAX:000664, The Jackson Laboratory) and were bred in house within a University of Florida vivarium. *drd1*-Cre (B6.FVB(Cg)-Tg(Drd1-cre)EY262Gsat/Mmucd, RRID:MMRRC_030989-UCD), *drd2*-Cre (B6.FVB(Cg)-Tg(Drd2-cre)ER44Gsat/Mmucd, RRID:MMRRC_032108-UCD), and *a2a*- Cre (B6.FVB(Cg)-Tg(Adora2a-cre)KG139Gsat/Mmucd, RRID:MMRRC_036158-UCD) mice were obtained from the UC Davis Mutant Mouse Regional Resource Center. Ai9 TdTomato Cre reporter mice (B6.Cg-Gt(ROSA)26Sor^tm9(CAG-tdTomato)Hze^/J; RRID:IMSR_JAX:007909, (Madisen et al., 2010)) were obtained from the Jackson Laboratory.

### Viral vectors

rgAAV.hSyn.HI.eGFP-Cre.WPRE.SV40 (Addgene #105540-AAVrg, 7×10^12^ vg/ml), Ef1α.DIO.Synaptophysin-mRuby and Ef1α.FLEX.Synaptophysin.GFP (both generous gifts from Dr Marc Fuccillo, University of Pennsylvania) (Herman et al., 2016), and AAV.hSyn.FLEx.mGFP-2A-Synaptophysin.mRuby (Addgene #71760-AAV1, 9.8×10¹² vg/mL) were used for tracing. AAV.Ef1a.DIO.hChR2(E123T/T159C)-EYFP (Addgene #35509-AAV5, 1x10^12^ vg/ml vg/ml) was used for patch-clamp recording and for optogenetic stimulation during the optogenetic real time place preference/avoidance task. AAV.Ef1a.DIO.EYFP (Addgene #27056-AAV5, 1×10^12^ vg/ml) was used as a control virus for the optogenetic real time place preference task. rgAAV.hSyn.DIO.hM4D(Gi)-mCherry (Addgene #44362-AAVrg, 1.2×10^13^ vg/ml) and rgAAV.hSyn.DIO.mCherry (50459-AAVrg, 1.8×10^13^ vg/ml) were used for chemogenetic inhibition.

### Surgical procedures

For all surgical procedures, mice were anesthetized with 2%–4% isoflurane (IsoFlo, Patterson Veterinary, Greeley, CO) in 1 L/min O_2_, and head fixed in a stereotaxic apparatus while their body temperature was maintained using a 38°C water bath heating pad. The scalp was shaved and cleaned with betadine and 70% ethanol. Following subcutaneous (s.c.) administration of Meloxicam (20 mg/kg) analgesia and local administration of the anesthetic lidocaine (lidocaine, 3 mg/kg, s.c., Patterson Veterinary) to the scalp, a small midline cranial incision was made.

For viral injections, craniotomies were made above the target regions. A pulled glass micropipette containing the AAV was slowly inserted for injection. For TuS injections, 50nl of viral solution was injected bilaterally at the following coordinates: anteroposterior (AP) +1.4mm bregma, mediolateral (ML) ±1.2mm lateral midline, dorsoventral (DV) - 4.85mm from the brain surface. For NAc injections, 100nl of viral solution was delivered bilaterally (AP 1.5mm, ML ±1.0mm, DV −3.75mm). For BLA injections, 100nl of viral solution was delivered either unilaterally into the right hemisphere (AP −1.6mm, ML +3.25mm, DV −4.25mm) for Opto-RTPP/A and brain slice electrophysiology experiments, or bilaterally (AP −1.6mm, ML ±3.25mm, DV −4.25mm) for tracing experiments. All injections were performed at a rate of 2nl/second (s), with 20-40s intervals using a Nanoject III (Drummond Scientific). Following injection, at least 5min went by before slowly withdrawing the pipette from the brain. Craniotomies were then sealed with dental wax and the incision was closed with wound clips.

For cannula implantation, the skull was scrubbed with 3% H_2_O_2_ and covered with a thin layer of cyanoacrylate glue (Vetbond, 3M). Bilateral craniotomies were drilled over the BLA and 26-gauge(G) guide cannulae (#C315GMN/SPC, P1 Technologies) extending 3.5mm below pedestal were implanted at the coordinates AP −1.3mm, ML ±3.2mm. Cannulae were then lowered into the brain and secured to the skull with a small amount of Vetbond followed by dental cement, and dust caps with a 3.5mm projection wire (C315DCMN/SPC, P1 Technologies) were inserted.

For optical fiber implantation, following skull preparation for implantation as above, a craniotomy was made and drilled above the ventral striatum on the right hemisphere. Fibers (300µM core diameter, 0.39NA, 6.0mm length) for optogenetic stimulation were lowered into the NAc (AP 1.4mm, ML 1.0mm, DV-3.85mm) or the TuS (AP 1.5mm, ML 1.2mm, DV −4.9mm). The fiber was secured with Vetbond followed by dental cement as described for the cannulae implantation.

Following surgery, mice were allowed to recover on a heating pad until ambulatory, and were given immediate *ad libitum* access to food and water. Meloxicam analgesic (20mg/kg, s.c.) was administered for at least 3 days following surgery. Mice will indwelling cranial implants were single housed and given 7 to 14 days after surgery to recover before the being acclimated to behavioral procedures.

### Histology

#### Immunohistochemistry

Mice were anesthetized with Fatalplus (150mg/kg; Vortech Pharmaceutical Ltd, Dearborn, MI) and transcardially perfused with cold 0.9% NaCl (Physiological Saline), followed by cold 10% phosphate buffered formalin (#SF100-4, Thermo Fisher Scientific) for fixation. Brains were collected and further fixed and cryoprotected in a 30% sucrose/10% formalin solution for 72hr at 4 °C. Serial 40μm thick coronal sections were collected using a sliding microtome (Leica) and were stored at 4 °C in a solution of Tris-buffered saline (TBS) with 0.03% sodium azide.

Sections from *drd1*- or *drd2*-Cre mice injected with Cre-dependent retrograde mCherry AAV underwent antigen retrieval in citrate buffer (pH 6.0) for 30mins at 80°C. After being rinsed with tris buffered saline (TBS; 0.242% Tris base, 2.924% sodium chloride, pH=7.4 ± 0.2) and diluting buffer (2% bovine serum albumin (Sigma Aldrich), 0.9% sodium chloride (Sigma Aldrich), 0.4% Triton-X 100 (Sigma Aldrich), and 1% normal goat serum (Sigma Aldrich) in TBS), samples were blocked in 20% normal donkey serum solution, then incubated in the primary antibody overnight at 4°C. Sections were then incubated in the secondary antibody at room temperature and washed with TBS prior to slide-mounting with DAPI Fluoromount-G® mounting medium (SouthernBiotech, catalog #0100-20). Primary antibodies included rabbit anti-DsRed (Takara Bio, catalog #632496, 1:1000) and chicken anti-NeuN/FOX3 (EnCor, catalog *#*CPCA-FOX3, 1:1000). Secondary antibodies included anti-chicken Alexa Fluor 488, anti-rabbit Alexa Fluor 680 (both from Invitrogen, 1:1000 dilution).

#### Imaging and quantification

Brain regions were identified using the mouse brain atlas (Paxinos and Franklin, 2000). Images were acquired using a Nikon Eclipse Ti2e fluorescent microscope. For quantification of the number of *drd1*+ and *drd2*+ TuS and NAc projecting BLA neurons, at least three BLA sections from three mice of each genotype and injection site were acquired spanning from −1.10mm to −2.10mm posterior to Bregma. Images were acquired at 20x magnification across both hemispheres and Z-stacked every 4µm. For quantification, regions of interest (ROIs) were drawn around the areas of interest (LA, BA). Images were preprocessed to remove background and to enhance local contrast, a rolling ball algorithm was applied to remove background, and images underwent Gaussian smoothing and Laplace sharpening. A semi-automated thresholding and counting algorithm created within NIS elements (Nikon) software was used to identify cells within selected ROIs, allowing for unbiased estimation of cell numbers. Cells were identified based on fluorescence intensity (via threshold) and diameter.

For quantification of *drd1+* and *a2a+* BLA to ventral striatum synaptophysin puncta within the ventral striatum, at least three sections from three mice of each genotype were acquired spanning from 1.7mm to 0.6mm anterior to Bregma. Images were acquired at 20x magnification for the hemisphere ipsilateral to the injection site, and Z-stacked every 0.9µm. For quantification, ROIs were drawn around the areas of interest (TuS, NAcC, NAcSh, PCX). Images were preprocessed to remove the average background. A semi-automated thresholding and counting algorithm created within NIS elements software was used to identify fluorescent puncta within selected ROIs, allowing for unbiased estimation of the number of fluorescent puncta. Puncta were identified based on fluorescence intensity (via threshold) and diameter.

#### Brain slice electrophysiology

Whole-cell patch-clamp recordings were performed in *ex vivo* brain slices from d*rd1*- Cre;Ai9 or a*2a*-Cre:Ai9 mice, in which tdTomato expression was directed within cells expressing either *drd1* or *drd2*, respectively. A Cre dependent AAV encoding for ChR2 (AAV-Ef1a-DIO hChR2(E123T/T159C)-EYFP) was injected bilaterally into the BLA of *drd1*-Cre:Ai9 or *a2a*-Cre:Ai9 mice, 2–3 months of age. After waiting a minimum of one month to allow for ample AAV expression, acute brain slices were prepared as follows.

Mice were deeply anesthetized with intraperitoneal injection of ketamine-xylazine (200- 15 mg/kg body weight) and decapitated. The cranium was dissected and the brain was immediately removed and placed in ice-cold HEPES buffered cutting solution containing (in mM): 92 N-methyl-d-glucamine, 2.5 KCl, 1.2 NaH2PO4, 30 NaHCO3, 20 HEPES, 25 glucose, 5 sodium l-ascorbate, 2 thiourea, 3 sodium pyruvate, 10 MgSO4 and 0.5 CaCl2 (osmolality ∼300 mOsm and pH ∼7.4, bubbled with 95% O2 and 5% CO2). Coronal brain slice (180-200µM) containing the OT were cut using a Leica VT 1200S vibratome. Brain slices were incubated in oxygenated artificial cerebrospinal fluid (ACSF) containing (in mM): 126 NaCl, 2.5 KCl, 2.4 CaCl2, 1.2 MgSO4, 1.4 NaH2PO4, 11 glucose, 25 NaHCO3 and 0.6 sodium L-ascorbate (osmolality ∼300mOsm and pH ∼7.4, bubbled with 95% O2 and 5% CO2) for 1hr at 31°C and at room temperature thereafter. Slices were transferred to the recording chamber for whole-cell patch-clamp recordings and continuously perfused with oxygenated ACSF. 4- Aminopyridine (4-AP; 200µM) was added to enhance optically evoked synaptic release in ChR2+ axonal terminals. Fluorescent D1-/A2A-tdTomato+ cells in OT were visualized with a 40 X water-immersion objective under an Olympus BX51WI upright microscope equipped with epifluorescence. Electrophysiological recordings were controlled by an EPC-10 amplifier combined with Pulse Software (HEKA Electronic) and analyzed using pulse and Clampfit (Axon instruments). Whole-cell patch-clamp recordings were made in both current and voltage-clamp mode. Patch pipettes were pulled from thin-wall borosilicate glass-capillary tubing (WPI, Sarasota, FL, USA) on a Flaming/Brown puller (P-97; Sutter Instruments Co., Novato, CA, USA). The tip resistance of the electrode was 5–8MΩ. The pipette solution contained the following (in mM): 120 K-gluconate, 10 NaCl, 1 CaCl2, 10 EGTA, 10 HEPES, 5 Mg-ATP, 0.5 Na-GTP, and 10 phosphocreatine.

To activate ChR2 in the OT slices, blue light (pE-300ultra, CoolLED, ∼25mW) was delivered through the same 40X objective. Pharmacological reagents including tetrodotoxin (TTX) citrate (Abcam), 6-cyano-7-nitroquinoxaline-2,3-dione (CNQX), d,l-2- amino-5-phosphonopentanoic acid (AP5), and 4-Aminopyridine (4-AP) (Sigma-Aldrich) were bath perfused during recording.

#### in vivo DREADD-based chemogenetic inhibition

For DREADD-based chemogenetic inhibition of Gi coupled inhibitory DREADD receptor (hM4Di) expressing neurons, *drd1*+ and *drd2*+ mice were injected with rgAAV.hSyn.DIO.hM4D(Gi)-mCherry (1.2x10^13^vg/ml, 100nl/hemisphere in NAc, 50nl/hemisphere in TuS, catalog #44362-AAVrg, Addgene) or rgAAV.hSyn.DIO.mCherry (1.8x10^13^vg/ml, 100nl/hemisphere in NAc, 50nl/hemisphere in TuS, catalog #50459-AAVrg, Addgene) as control. All mice were implanted 1-2 weeks later with bilateral intracranial guide cannulae (Protech International, Inc, catalog #8IC315GMNSPC, 26ga) extending 3.5mm beyond the pedestal, for direct administration of either the DREADD ligand J60 (Bonaventura et al., 2019) or vehicle into the BLA. Dust caps without a projection wire (Protech International, Inc, catalog #8IC315DCMNSP) were inserted immediately following surgery, and mice were given 1-2 weeks to recover.

Prior to behavior, mice underwent 2 days of handling in which the dummy cannulae were removed and replaced. On the habituation behavior day, mice received a “mock” infusion, wherein the internal cannulae (Protech International, Inc, catalog #8IC315MNSPC, 5.75mm projection, 33ga) connected to tubing from a 1µL Hamilton Syringe (Hamilton, catalog #86211) were inserted into the guide cannulae, and the Harvard Apparatus 22 Syringe Pump (catalog #PY2 55-2222) was turned on for 2 min to simulate the noise of the infusion. This mock infusion occurred 30 min prior to being placed in the plethysmograph for the Pavlovian fear learning behavioral paradigm, and occurred on a separate day from the spontaneous odor attraction/avoidance assay. On the learning day (Day 2) of the Pavlovian fear learning paradigm, and on the day of the spontaneous odor attraction/avoidance assay, mice were once again tethered to the Hamilton syringe, but this time received an infusion of 100nL of either 10nM J60 or vehicle at a rate of 50nl/min, 30 min prior to the start of the behavioral task.

## Behavioral Tasks

### Odor-shock Pavlovian fear learning

We used a whole-body plethysmography chamber (Data Sciences International, St. Paul, MN) that was adapted for the infusion of a neutral odor and the administration of a mild foot shock for an odor-shock Pavlovian fear learning test, as originally developed for use in rats (Hegoburu et al., 2011). We constructed an air-dilution olfactometer (Gadziola et al., 2015; Johnson et al., 2020) and used custom code in Synapse (Tucker Davis Technologies) to control the delivery of an otherwise neutral odor, isopentyl acetate (1 torr in liquid state; Sigma Aldrich), at a flow rate of 1 L/min (20s) which co-terminated with the presentation of a mild foot shock (0.5mA for 1s). Respiratory transients were detected using a Data Sciences pressure transducer, gain amplified 100 X (Cygnus Technology Inc), and digitized (0.1-20Hz) at 300Hz in Synapse. Positive pressure of clean room air was continuously applied to the chamber using a stable- output air pump (Tetra Whisper). Following each stimulus trial, odor-vaporized air was exhausted from the plethysmograph through an outlet at the chamber’s ceiling.

Mice were acclimated to handling in the behavioral room for two days prior to entering the plethysmograph. Mice were then acclimated to the plethysmograph by undergoing a session in which no odors or shock were delivered, but the associated sounds were present (**Supplementary Fig 4**). Twenty-four hr later on the acquisition day, mice were allowed to acclimate to the plethysmograph for a 4-minute (min) period and were then presented with 10 trials of 20s odor delivery co-terminating with an odor-paired 1s foot shock (0.5mA) with an inter-trial interval (ITI) of 180s. For the unpaired fear conditioning task, the foot shock was presented pseudorandomly in the ITI (90s after the foot shock). For the odor only control mice, the 10 trials consisted of only 20s odor delivery without the administration of the foot shock. The shocked mice received no odor delivery during the trials, but received a foot shock either at the end of the trial (trial shock group) or pseudorandomly in the ITI (ITI-shock group). Mice were then returned to their home cage. Twenty-four hr later on the retrieval day, the odor was presented for 10 trials without the foot shock for all groups receiving odor (paired, unpaired, and odor only groups). Mice who did not previously receive the odor underwent the 10 trials without odor delivery or foot shock. Mobility behavior was recorded throughout the entire fear conditioning task using two digital cameras (Microsoft, 10Hz frame rate), and was scored in 0.4s bins during the 19s of odor presentation prior to shock using ezTrack (Pennington et al., 2019) to identify periods of physical immobility. Respiration digitized from the pressure transducer was imported into MATLAB and a MATLAB script was used to calculate fast-fourier transform (FFT) power spectra of the respiratory signal during odor (excluding the 1s when the shock co-occurred) as compared to pre-odor (see **Supplementary Fig 4**).

### Spontaneous odor attraction or avoidance

To test the spontaneous attraction or avoidance towards odors, a 30.48 × 30.48 × 30.48 cm (length × width × height) dark acrylic chamber was divided into two equal sides by a transparent acrylic plate with a tunnel in the bottom center to allow mice to pass through (**Fig 6**). An infrared video camera was placed above the chamber to record activity of the mouse in each chamber (12Hz frame rate).

Cotton swabs laced with peanut oil (diluted in mineral oil, 1:12.5), an appetitive odor, were placed in a perforated microcentrifuge tubes to prevent touching or tasting the stimulus while still allowing the release of volatiles. Tubes containing this appetitive odor were placed on one side of the chamber, while perforated microcentrifuge tubes containing cotton swabs laced with 2-Methyl-2-thiazoline (2MT, 97%, Fisher Scientific, diluted in mineral oil, 1:50), a component of fox feces, were placed on the opposite side. These dilutions were selected to achieve a comparable intensity of odor from each tube.

Mice were handled for two days prior to the behavioral assay to acclimate the mice to experimenter handling, and mice received a mock infusion to acclimate the mice to the sound of the infusion pump. On the day of the behavioral assay, mice received an infusion of either J60 or vehicle 30min prior to being placed in the center of the chamber, with the odors arranged on opposite sides. Mice were allowed to explore the appetitive peanut oil and aversive fox urine sides for 10min. All testing was performed in a dark room with a single dim light to illuminate subjects, and infrared video recordings were used to assess the amount of time spent in each side, after subtracting out the middle third of the apparatus – i.e. location of the tunnel. Analyses were performed in ezTrack (Pennington et al., 2019).

### Optogenetic real time place preference or aversion test (Opto-RTPP/A)

Mice were gently handled and acclimated to the behavior room the day prior to the opto- RTPP/A test. Prior to starting the opto-RTPP/A test, mice were gently scruffed, the dust cap was removed, and the mice were tethered to a 400µm, 0.57NA fiber (Thorlabs, catalog #M58L01) and placed in a 15.24 × 40.64 × 27.94 cm (length × width × height) apparatus divided into three chambers. This fiber was connected to an LED (Doric, 465nm) through a rotary joint connected to a 400µm, 0.39NA patch cable. Mice were placed in the center of a three-chamber apparatus and allowed to explore for 30min. An infrared video camera was placed above the chamber to record activity of the mouse in each chamber (12Hz frame rate). When mice entered into one of the three chambers, and subsequently broke the infrared beam path, light stimulation (465nm, 15ms pulse width, 40Hz) was initiated and continuously delivered until mice left the chamber and ceased breaking the infrared beams (controlled by an Arduino). At the end of the 30min, mice were gently restrained and the tether was removed, following which the mice were returned to their home cage. The mice were euthanized and perfused the same day, and brains were collected for histological verification of virus injection and optic fiber placement. Analyses were performed in ezTrack (Pennington et al., 2019) to quantify the time spent in each chamber and to generate maps of physical space for illustration purposes.

---

**Supplemental Figure 1.**
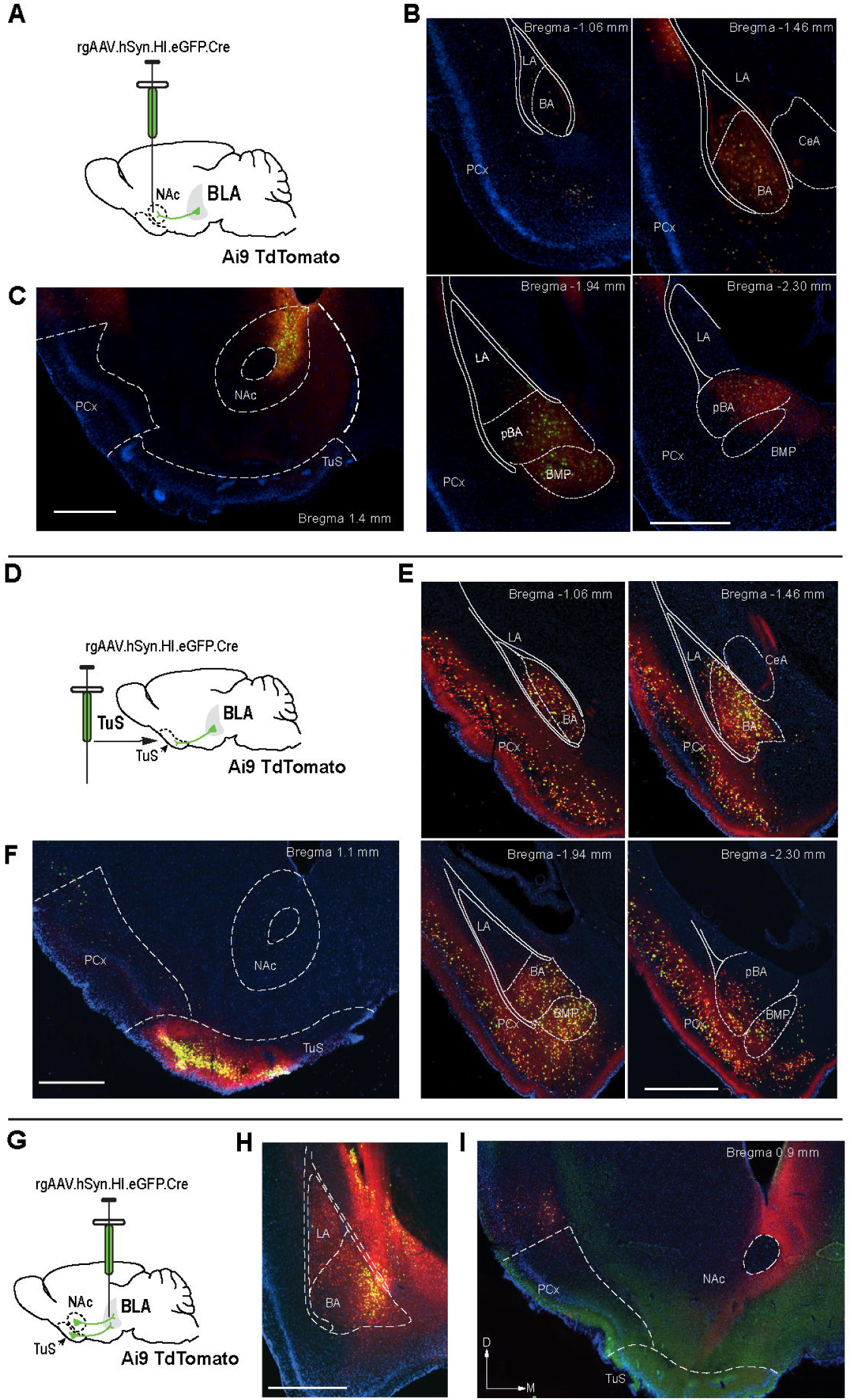

**Supplemental Figure 2.**
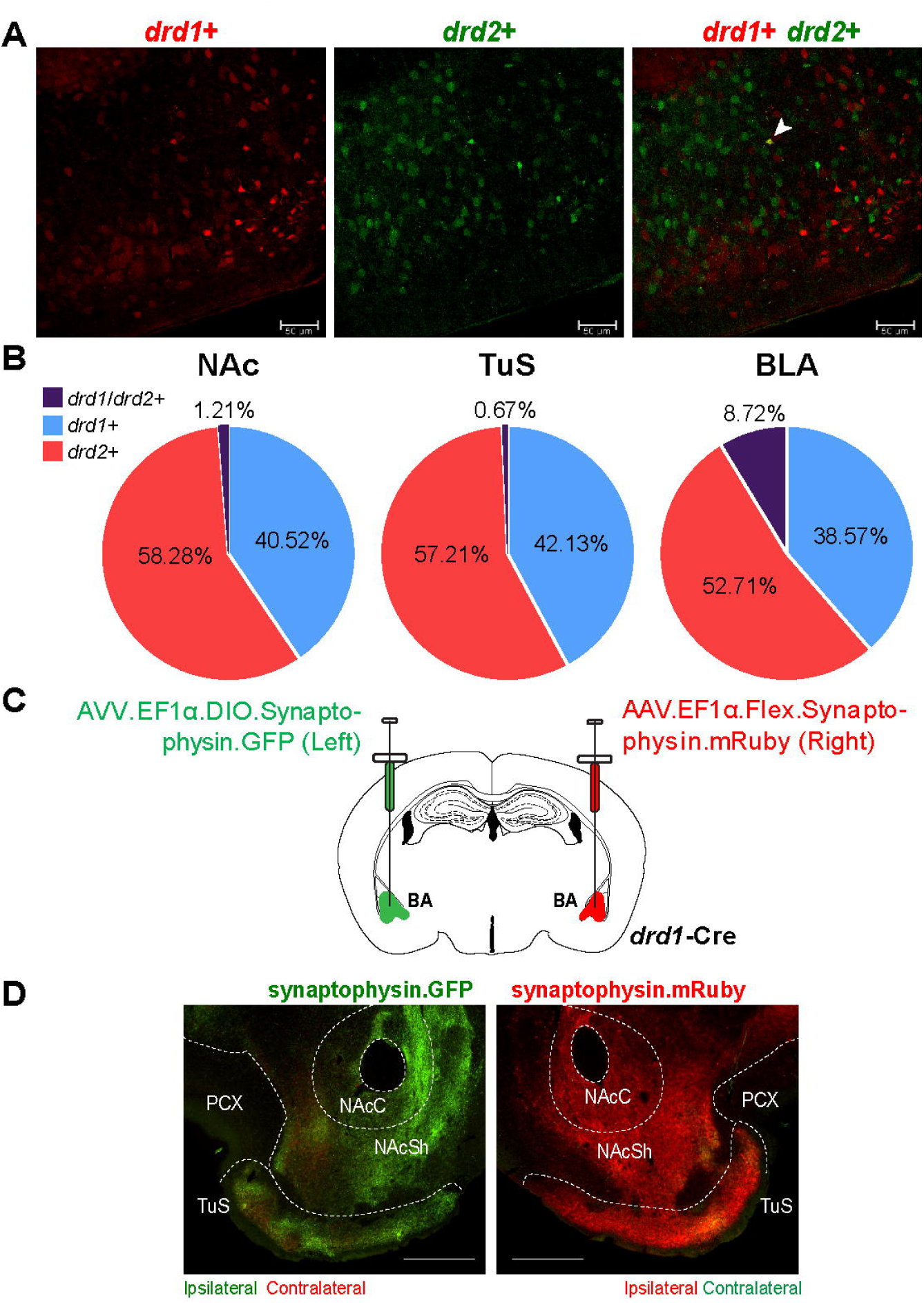

**Supplemental Figure 3.**
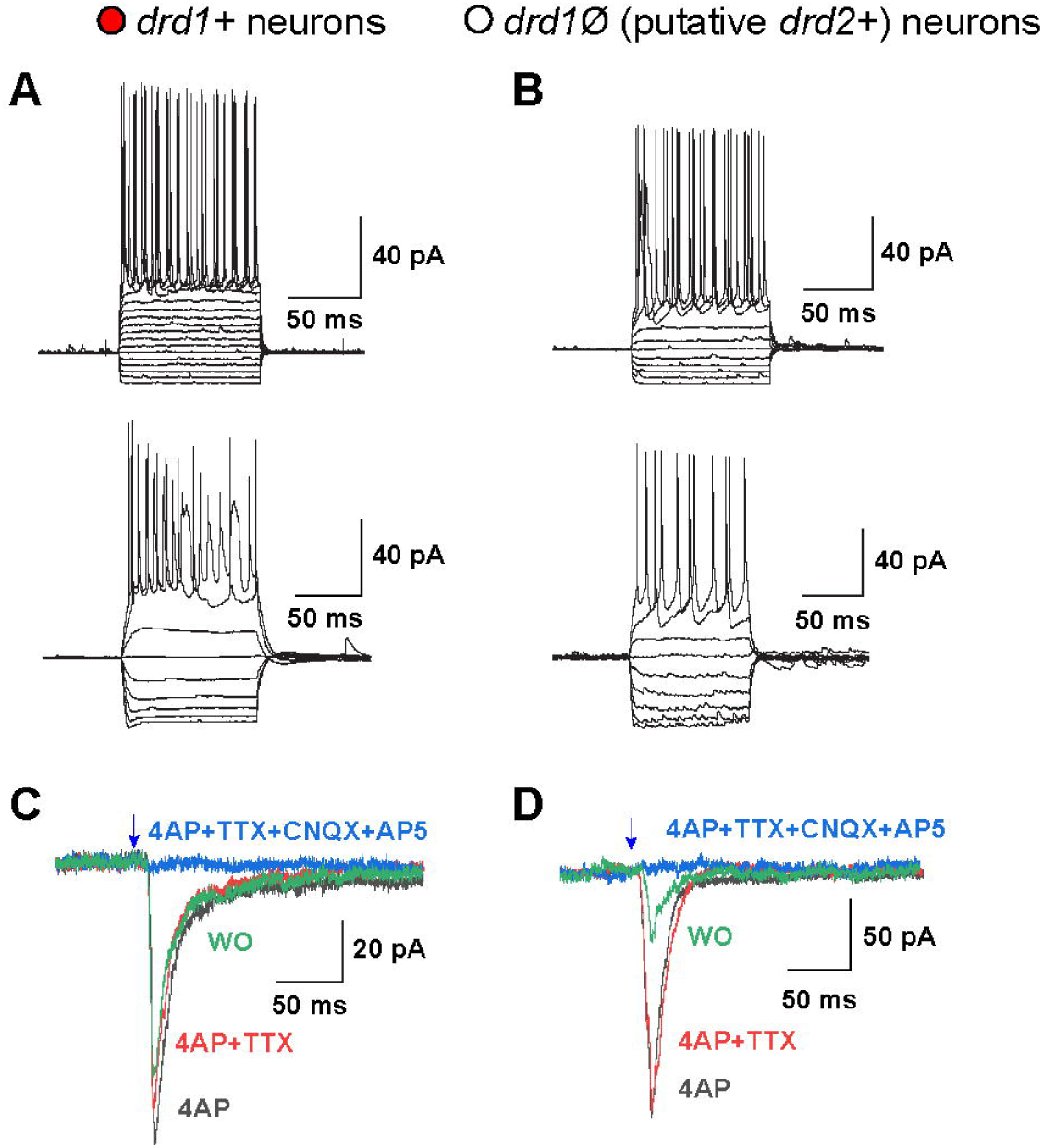

**Supplemental Figure 4.**
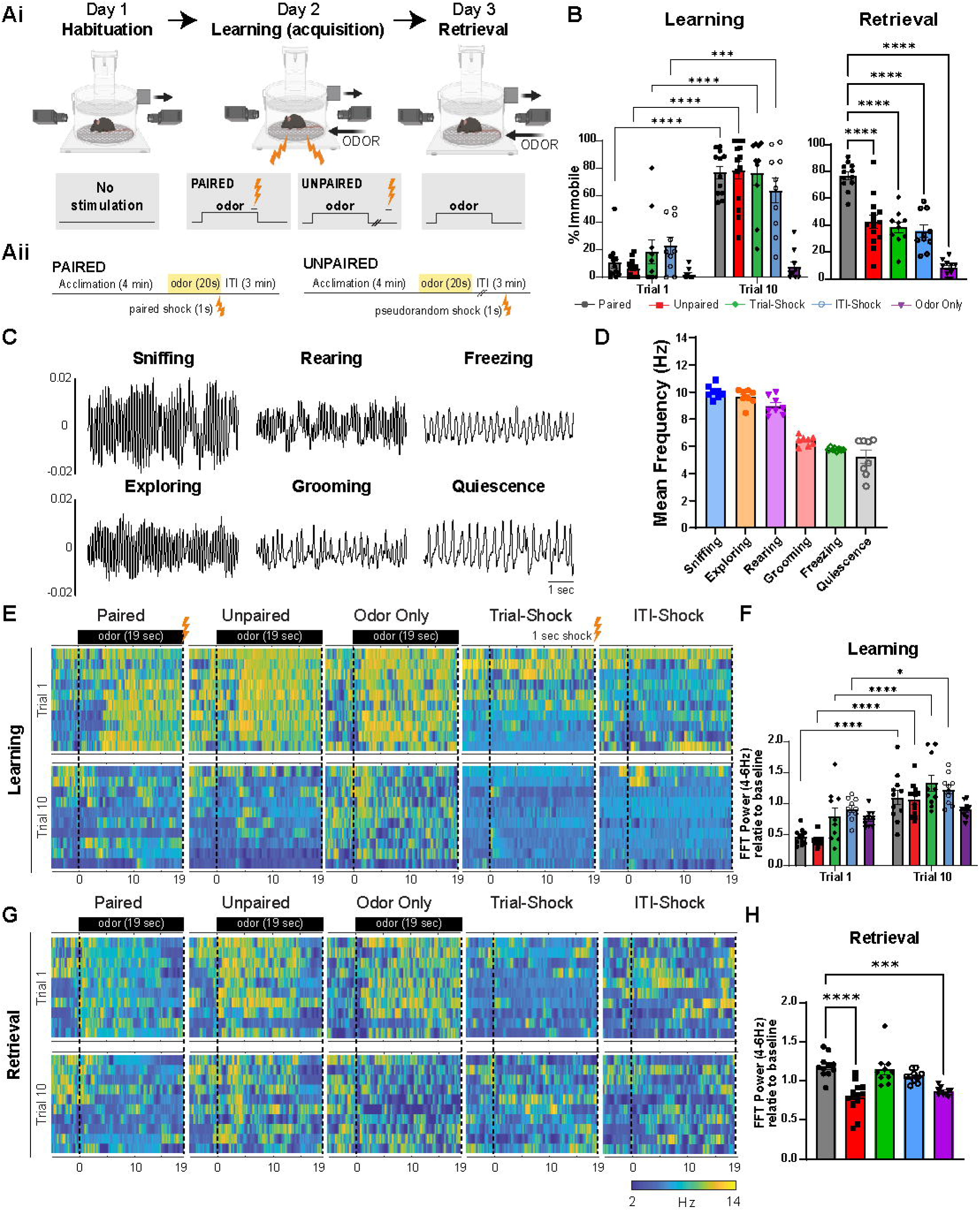

**Supplemental Figure 5.**
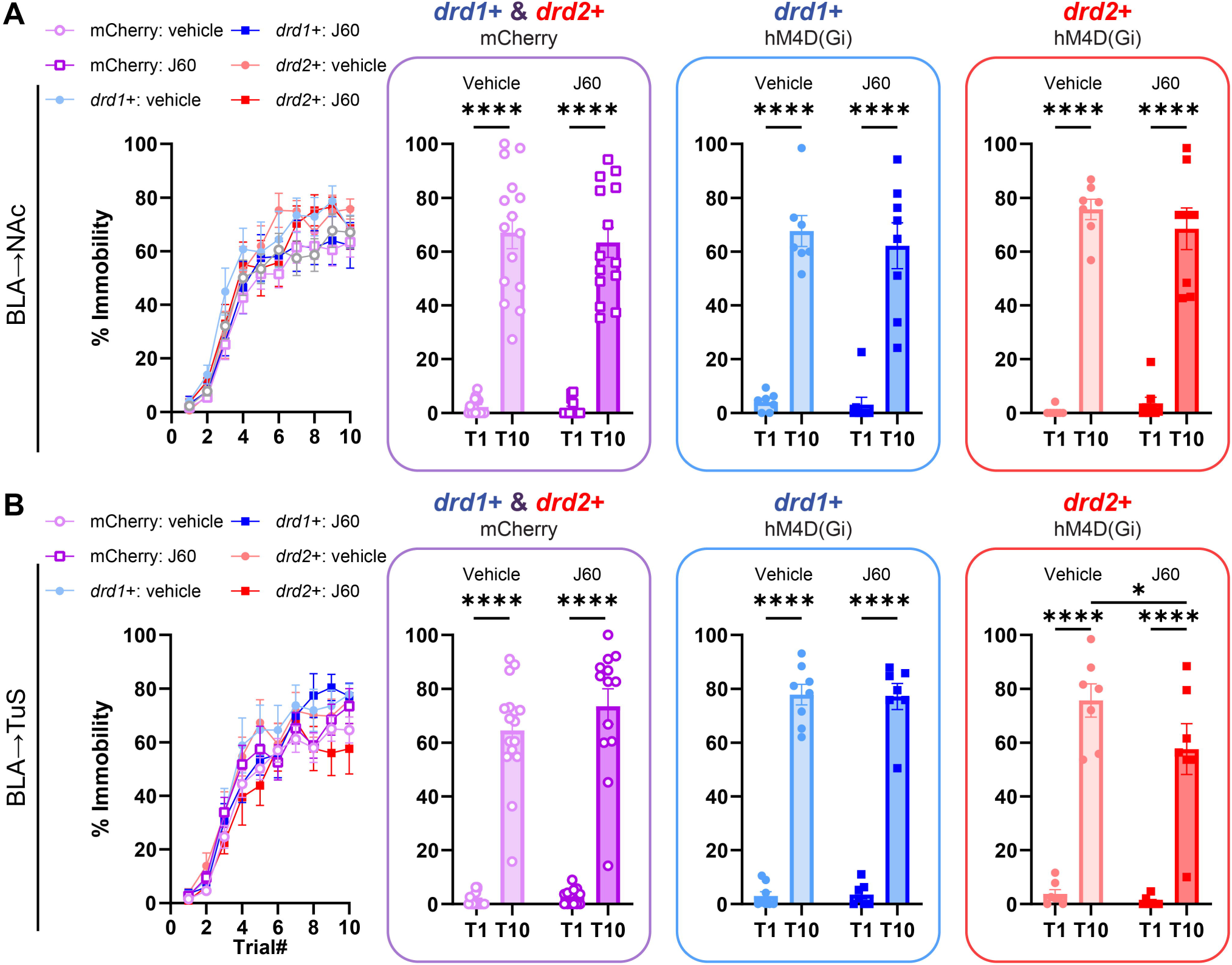

## Notes

### Competing Interest Statement

The authors have declared no competing interest.

## References

1. Ambroggi F, Ishikawa A, Fields HL, Nicola SM (2008) Basolateral Amygdala Neurons Facilitate Reward-Seeking Behavior by Exciting Nucleus Accumbens Neurons. Neuron 59:648–661.

2. Berglind WJ, Case JM, Parker MP, Fuchs RA, See RE (2006) Dopamine D1 or D2 receptor antagonism within the basolateral amygdala differentially alters the acquisition of cocaine-cue associations necessary for cue-induced reinstatement of cocaine-seeking. Neuroscience 137:699–706.

3. Best AR, Wilson DA (2003) A postnatal sensitive period for plasticity of cortical afferents but not cortical association fibers in rat piriform cortex. Brain Res 961:81–87 Available at: http://www.ncbi.nlm.nih.gov/entrez/query.fcgi?cmd=Retrieve&db=PubMed&dopt=Citation&list_uids=12535779.

4. Beyeler A, Namburi P, Glober GF, Simonnet C, Calhoon GG, Conyers GF, Luck R, Wildes CP, Tye KM (2016) Divergent Routing of Positive and Negative Information from the Amygdala during Memory Retrieval. Neuron 90:348–361 Available at: https://www.sciencedirect.com/science/article/pii/S0896627316001835.

5. Blanchard DC, Blanchard RJ (1972) Innate and conditioned reactions to threat in rats with amygdaloid lesions. J Comp Physiol Psychol 81:281–290.

6. Boiten FA (1998) The effects of emotional behaviour on components of the respiratory cycle. Biol Psychol 49.

7. Bonaventura J et al. (2019) High-potency ligands for DREADD imaging and activation in rodents and monkeys. Nat Commun 10:4627 Available at: 10.1038/s41467-019-12236-z.

8. Bristol AS, Sutton MA, Carew TJ (2004) Neural circuit of tail-elicited siphon withdrawal in Aplysia. I. Differential lateralization of sensitization and dishabituation. J Neurophysiol 91:666–677 Available at: http://www.ncbi.nlm.nih.gov/entrez/query.fcgi?cmd=Retrieve&db=PubMed&dopt=Citation&list_uids=13679401.

9. Cahill L, McGaugh JL (1990) Amygdaloid complex lesions differentially affect retention of tasks using appetitive and aversive reinforcement. Behav Neurosci 104:532– 543.

10. Cardinal RN, Parkinson JA, Hall J, Everitt BJ (2002) Emotion and motivation: the role of the amygdala, ventral striatum, and prefrontal cortex. Neurosci Biobehav Rev 26:321–352 Available at: http://www.ncbi.nlm.nih.gov/entrez/query.fcgi?cmd=Retrieve&db=PubMed&dopt=Citation&listuids=12034134.

11. Cousens G, Otto T (1998) Both pre- and posttraining excitotoxic lesions of the basolateral amygdala abolish the expression of olfactory and contextual fear conditioning. Behav Neurosci 112:1092–1103.

12. de Oliveira AR, Reimer AE, Macedo CEA de, Carvalho MC de, Silva MA de S, Brandão ML (2011) Conditioned fear is modulated by D2 receptor pathway connecting the ventral tegmental area and basolateral amygdala. Neurobiol Learn Mem 95:37–45 Available at: https://www.sciencedirect.com/science/article/pii/S1074742710001723.

13. de Souza Caetano KA, de Oliveira AR, Brandão ML (2013) Dopamine D2 receptors modulate the expression of contextual conditioned fear: role of the ventral tegmental area and the basolateral amygdala. Behav Pharmacol 24 Available at: https://journals.lww.com/behaviouralpharm/Fulltext/2013/08000/Dopamine_D2_receptors_modulate_the_expression_of.4.aspx.

14. Fadok JP, Dickerson TMK, Palmiter RD (2009) Dopamine is necessary for cue- dependent fear conditioning. J Neurosci.

15. Gadziola MA, Tylicki KA, Christian DL, Wesson DW (2015) The Olfactory Tubercle Encodes Odor Valence in Behaving Mice. J Neurosci 35:4515–4527 Available at: http://www.jneurosci.org/content/35/11/4515.abstract.

16. Gong S, Doughty M, Harbaugh CR, Cummins A, Hatten ME, Heintz N, Gerfen CR (2007) Targeting Cre Recombinase to Specific Neuron Populations with Bacterial Artificial Chromosome Constructs. J Neurosci 27:9817 LP – 9823 Available at: http://www.jneurosci.org/content/27/37/9817.abstract.

17. Greba Q, Gifkins A, Kokkinidis L (2001) Inhibition of amygdaloid dopamine D2 receptors impairs emotional learning measured with fear-potentiated startle. Brain Res 899:218–226 Available at: https://www.sciencedirect.com/science/article/pii/S0006899301022430.

18. Guarraci FA, Frohardt RJ, Falls WA, Kapp BS (2000) The effects of intra-amygdaloid infusions of a D₂ dopamine receptor antagonist on Pavlovian fear conditioning. Behav Neurosci 114:647–651.

19. Guarraci FA, Frohardt RJ, Young SL, Kapp BS (1999) A Functional Role for Dopamine Transmission in the Amygdala during Condtioned Fear. Ann N Y Acad Sci 877:732–736 Available at: 10.1111/j.1749-6632.1999.tb09312.x.

20. Hatfield T, Han J-S, Conley M, Gallagher M, Holland P (1996) Neurotoxic Lesions of Basolateral, But Not Central, Amygdala Interfere with Pavlovian Second-Order Conditioning and Reinforcer Devaluation Effects. J Neurosci 16:5256 LP – 5265 Available at: http://www.jneurosci.org/content/16/16/5256.abstract.

21. Hegoburu C, Shionoya K, Garcia S, Messaoudi B, Thévenet M, Mouly A-M (2011) The RUB Cage: Respiration–Ultrasonic Vocalizations–Behavior Acquisition Setup for Assessing Emotional Memory in Rats. Front Behav Neurosci 5 Available at: http://www.frontiersin.org/Journal/Abstract.aspx?s=99&name=behavioral_neuroscience&ART DOI=10.3389/fnbeh.2011.00025.

22. Herman AM, Ortiz-Guzman J, Kochukov M, Herman I, Quast KB, Patel JM, Tepe B, Carlson JC, Ung K, Selever J, Tong Q, Arenkiel BR (2016) A cholinergic basal forebrain feeding circuit modulates appetite suppression. Nature 538:253–256 Available at: 10.1038/nature19789.

23. Hochgerner H, Singh S, Tibi M, Lin Z, Skarbianskis N, Admati I, Ophir O, Reinhardt N, Netser S, Wagner S, Zeisel A (2023) Neuronal types in the mouse amygdala and their transcriptional response to fear conditioning. Nat Neurosci 26:2237–2249 Available at: 10.1038/s41593-023-01469-3.

24. Homma I, Masaoka Y (2008) Breathing rhythms and emotions. Exp Physiol 93.

25. Horvitz JC (2000) Mesolimbocortical and nigrostriatal dopamine responses to salient non-reward events. Neuroscience 96:651–656 Available at: https://www.sciencedirect.com/science/article/pii/S0306452200000191.

26. Janak PH, Tye KM (2015) From circuits to behaviour in the amygdala. Nature 517:284 Available at: 10.1038/nature14188.

27. Johnson ME, Bergkvist L, Mercado G, Stetzik L, Meyerdirk L, Wolfrum E, Madaj Z, Brundin P, Wesson DW (2020) Deficits in olfactory sensitivity in a mouse model of Parkinson’s disease revealed by plethysmography of odor-evoked sniffing. Sci Rep 10:9242 Available at: https://www.nature.com/articles/s41598-020-66201-8.

28. Jones S V, Choi DC, Davis M, Ressler KJ (2008) Learning-Dependent Structural Plasticity in the Adult Olfactory Pathway. J Neurosci 28:13106 LP – 13111 Available at: http://www.jneurosci.org/content/28/49/13106.abstract.

29. Kim ES, Lattal KM (2019) Context-Dependent and Context-Independent Effects of D1 Receptor Antagonism in the Basolateral and Central Amygdala during Cocaine Self-Administration. eNeuro 6.

30. Kim J, Pignatelli M, Xu S, Itohara S, Tonegawa S (2016) Antagonistic negative and positive neurons of the basolateral amygdala. Nat Neurosci 19:1636–1646 Available at: 10.1038/nn.4414.

31. Kim J, Zhang X, Muralidhar S, LeBlanc SA, Tonegawa S (2017) Basolateral to Central Amygdala Neural Circuits for Appetitive Behaviors. Neuron 93:1464–1479.e5 Available at: http://www.sciencedirect.com/science/article/pii/S0896627317301423.

32. Klüver H, Bucy PC (1939) Preliminary analysis of functions of the temporal lobes in monkeys. Arch Neurol Psychiatry 42:979–1000.

33. Kröner S, Rosenkranz JA, Grace AA, Barrionuevo G (2005) Dopamine Modulates Excitability of Basolateral Amygdala Neurons In Vitro. J Neurophysiol 93:1598– 1610 Available at: 10.1152/jn.00843.2004.

34. Lamont EW, Kokkinidis L (1998) Infusion of the dopamine D1 receptor antagonist SCH 23390 into the amygdala blocks fear expression in a potentiated startle paradigm. Brain Res 795:128–136 Available at: https://www.sciencedirect.com/science/article/pii/S0006899398002819.

35. LeDoux J (2003) The emotional brain, fear, and the amygdala. Cell Mol Neurobiol 23:727–738 Available at: http://www.ncbi.nlm.nih.gov/entrez/query.fcgi?cmd=Retrieve&db=PubMed&dopt=Ci tation&list_uids=14514027.

36. LeDoux JE (1992) Brain mechanisms of emotion and emotional learning. Curr Opin Neurobiol 2:191–197 Available at: http://www.ncbi.nlm.nih.gov/entrez/query.fcgi?cmd=Retrieve&db=PubMed&dopt=Ci tation&list_uids=1638153.

37. Li W, Howard JD, Parrish TB, Gottfried JA (2008) Aversive learning enhances perceptual and cortical discrimination of indiscriminable odor cues. Science (80-) 319:1842–1845 Available at: http://www.ncbi.nlm.nih.gov/entrez/query.fcgi?cmd=Retrieve&db=PubMed&dopt=Citation&list_uids=18369149.

38. Madisen L, Zwingman TA, Sunkin SM, Oh SW, Zariwala HA, Gu H, Ng LL, Palmiter RD, Hawrylycz MJ, Jones AR, Lein ES, Zeng H (2010) A robust and high-throughput Cre reporting and characterization system for the whole mouse brain. Nat Neurosci 13:133–140 Available at: http://www.ncbi.nlm.nih.gov/pmc/articles/PMC2840225/.

39. Maren S, Aharonov G, Fanselow MS (1996) Retrograde abolition of conditional fear after excitotoxic lesions in the basolateral amygdala of rats: Absence of a temporal gradient. Behav Neurosci 110:718–726.

40. Maren S, Quirk GJ (2004) Neuronal signalling of fear memory. Nat Rev Neurosci 5.

41. Meador-Woodruff JH, Mansour A, Healy DJ, Kuehn R, Zhou QY, Bunzow JR, Akil H, Civelli O, Watson SJJ (1991) Comparison of the distributions of D1 and D2 dopamine receptor mRNAs in rat brain. Neuropsychopharmacol Off Publ Am Coll Neuropsychopharmacol 5:231–242.

42. Moberly AH, Schreck M, Bhattarai JP, Zweifel LS, Luo W, Ma M (2018) Olfactory inputs modulate respiration-related rhythmic activity in the prefrontal cortex and freezing behavior. Nat Commun 9:1528 Available at: 10.1038/s41467-018-03988-1.

43. Murata K, Kanno M, Ieki N, Mori K, Yamaguchi M (2015) Mapping of Learned Odor- Induced Motivated Behaviors in the Mouse Olfactory Tubercle. J Neurosci 35:10581–10599 Available at: http://www.jneurosci.org/content/35/29/10581.abstract.

44. Namburi P, Beyeler A, Yorozu S, Calhoon G, Halbert S, Wichmann R, Holden S, Mertens K, Anahtar M, Felix-Ortiz A, Wickersham I, Gray J, Tye K (2015) A circuit mechanism for differentiating positive and negative associations. Nature 520:675– 678 Available at: https://pubmed.ncbi.nlm.nih.gov/25925480/ [Accessed July 13, 2021].

45. Nishi A, Snyder GL, Greengard P (1997) Bidirectional Regulation of DARPP-32 Phosphorylation by Dopamine. J Neurosci 17:8147 LP – 8155 Available at: http://www.jneurosci.org/content/17/21/8147.abstract.

46. Ouimet CC, Miller PE, Hemmings HC, Walaas SI, Greengard P (1984) DARPP-32, a dopamine- and adenosine 3’:5’-monophosphate-regulated phosphoprotein enriched in dopamine-innervated brain regions. III. Immunocytochemical localization. J Neurosci 4:111–124.

47. Paré D, Quirk GJ, Ledoux JE (2004) New Vistas on Amygdala Networks in Conditioned Fear. J Neurophysiol 92:1–9 Available at: http://jn.physiology.org/content/92/1/1.abstract.

48. Paxinos G, Franklin K (2000) The Mouse Brain in Stereotaxic Coordinates, 2nd ed. San Diego: Academic Press.

49. Pennington ZT, Dong Z, Feng Y, Vetere LM, Page-Harley L, Shuman T, Cai DJ (2019) ezTrack: An open-source video analysis pipeline for the investigation of animal behavior. Sci Rep 9:19979 Available at: 10.1038/s41598-019-56408-9.

50. Richfield EK, Penney JB, Young AB (1989) Anatomical and affinity state comparisons between dopamine D1 and D2 receptors in the rat central nervous system. Neuroscience 30:767–777 Available at: https://www.sciencedirect.com/science/article/pii/0306452289901681.

51. Rogan MT, Staubli U V, LeDoux JE (1997) Fear conditioning induces associative long- term potentiation in the amygdala. Nature 390:604–607 Available at: http://www.ncbi.nlm.nih.gov/entrez/query.fcgi?cmd=Retrieve&db=PubMed&dopt=Citation&list_uids=9403688.

52. Schoenbaum G, Setlow B, Nugent SL, Saddoris MP, Gallagher M (2003) Lesions of orbitofrontal cortex and basolateral amygdala complex disrupt acquisition of odor- guided discriminations and reversals. Learn Mem 10:129–140.

53. Schultz W (2007) Multiple DA functions at different time courses. Annu Rev Neurosci 30:259.

54. Scibilia RJ, Lachowicz JE, Kilts CD (1992) Topographic nonoverlapping distribution of D1 and D2 dopamine receptors in the amygdaloid nuclear complex of the rat brain. Synapse 11:146–154.

55. See RE, Kruzich PJ, Grimm JW (2001) Dopamine, but not glutamate, receptor blockade in the basolateral amygdala attenuates conditioned reward in a rat model of relapse to cocaine-seeking behavior. Psychopharmacology (Berl) 154:301–310.

56. Setlow B, Gallagher M, Holland PC (2002) The basolateral complex of the amygdala is necessary for acquisition but not expression of CS motivational value in appetitive Pavlovian second-order conditioning. Eur J Neurosci 15:1841–1853 Available at: 10.1046/j.1460-9568.2002.02010.x.

57. Shionoya K, Hegoburu C, Brown B, Sullivan R, Doyère V, Mouly A-M (2013) It’s time to fear! Interval timing in odor fear conditioning in rats. Front Behav Neurosci 7 Available at: https://www.frontiersin.org/articles/10.3389/fnbeh.2013.00128.

58. Shuen JA, Chen M, Gloss B, Calakos N (2008) Drd1a-tdTomato BAC Transgenic Mice for Simultaneous Visualization of Medium Spiny Neurons in the Direct and Indirect Pathways of the Basal Ganglia. J Neurosci 28:2681–2685 Available at: http://www.jneurosci.org/cgi/doi/10.1523/JNEUROSCI.5492-07.2008.

59. Stevenson I, Ripley HS (1952) Variations in respiration and in respiratory symptoms during changes in emotion. Psychosom Med 14.

60. Stuber GD, Sparta DR, Stamatakis AM, van Leeuwen WA, Hardjoprajitno JE, Cho S, Tye KM, Kempadoo KA, Zhang F, Deisseroth K, Bonci A (2011) Excitatory transmission from the amygdala to nucleus accumbens facilitates reward seeking. Nature 475:377 Available at: 10.1038/nature10194.

61. Subramanian HH, Holstege G (2013) Stimulation of the midbrain periaqueductal gray modulates preinspiratory neurons in the ventrolateral medulla in the rat in vivo. J Comp Neurol 521:3083–3098 Available at: https://onlinelibrary.wiley.com/doi/10.1002/cne.23334.

62. Svenningsson P, Nishi A, Fisone G, JA G, AC N, Greengard P (2004) DARPP-32: an integrator of neurotransmission. Annu Rev Pharmacol Toxicol 44:269.

63. Walker P, Carrive P (2003) Role of ventrolateral periaqueductal gray neurons in the behavioral and cardiovascular responses to contextual conditioned fear and poststress recovery. Neuroscience 116:897–912 Available at: http://www.sciencedirect.com/science/article/pii/S0306452202007443.

64. Weiskrantz L (1956) Behavioral changes associated with ablation of the amygdaloid complex in monkeys. J Comp Physiol Psychol 49:381–391 Available at: https://psycnet.apa.org/journals/com/49/4/381 [Accessed July 15, 2021].

65. Wesson DW (2020) The Tubular Striatum. J Neurosci 40:7379–7386.

66. West EA, Carelli RM (2016) Nucleus Accumbens Core and Shell Differentially Encode Reward-Associated Cues after Reinforcer Devaluation. J Neurosci 36:1128–1139 Available at: http://www.jneurosci.org/content/36/4/1128.abstract.

67. White KA, Zhang Y-F, Zhang Z, Bhattarai JP, Moberly AH, in ‘t Zandt E, Peña-Bravo JI, Mi H, Jia X, Fuccillo M V., Xu F, Ma M, Wesson DW (2019) Glutamatergic neurons in the piriform cortex influence the activity of D1 and D2-type receptor expressing olfactory tubercle neurons. J Neurosci 38:9546–9559 Available at: http://www.ncbi.nlm.nih.gov/pubmed/31628176.

68. Wilson DA, Sullivan RM (2011) Cortical processing of odor objects. Neuron 72:506– 519.

69. Zhang X, Guan W, Yang T, Furlan A, Xiao X, Yu K, An X, Galbavy W, Ramakrishnan C, Deisseroth K, Ritola K, Hantman A, He M, Josh Huang Z, Li B (2021) Genetically identified amygdala–striatal circuits for valence-specific behaviors. Nat Neurosci 24:1586–1600 Available at: 10.1038/s41593-021-00927-0.

70. Zhang X, Kim J, Tonegawa S (2020a) Amygdala Reward Neurons Form and Store Fear Extinction Memory. Neuron 105:1077–1093.e7 Available at: https://www.sciencedirect.com/science/article/pii/S0896627319310918.

71. Zhang X, Kim J, Tonegawa S (2020b) Amygdala Reward Neurons Form and Store Fear Extinction Memory. Neuron 105:1077–1093.e7.

72. Zhang Z, Liu Q, Wen P, Zhang J, Rao X, Zhou Z, Zhang H, He X, Li J, Zhou Z, Xu X, Zhang X, Luo R, Lv G, Li H, Cao P, Wang L, Xu F (2017) Activation of the dopaminergic pathway from VTA to the medial olfactory tubercle generates odor- preference and reward. Elife 6:e25423.

