## Supplementary material for "Bidirectional modulation of negative emotional states by parallel genetically-distinct basolateral amygdala pathways to ventral striatum subregions": Supplmentary materials

^+^Indicates co-2^nd^ author

Shortened title: Basolateral amygdala circuitry to ventral striatum

**Supplemental Figures 1. BLA neurons directly innervating the ventral striatum. Related to Figures 1 and 2. (A)** Approach for identifying BLA inputs to the NAc in which a retrograde virus expressing Cre recombinase and GFP was unilaterally injected into the NAc of Ai9 TdTomato Cre-reporter mice. **(B)** Representative images from one mouse throughout the anterior-posterior extent of the BLA following rgAAV.hSyn.HI.eGFP.Cre injection in the NAc. **(C)** Example of the NAc injection site from the mouse in (B) illustrating confirming a region-specific injection. **(D)** Approach for identifying BLA inputs to the TuS. **(E)** Representative images from one mouse throughout the anterior-posterior extent of the BLA following rgAAV.hSyn.HI.eGFP.Cre injection in the TuS. **(F)** Example of the TuS injection site from the mouse in (E) illustrating confirming a region-specific injection. **(G)** Schematic of approach for identifying ventral striatum inputs to the BLA to determine if there is reciprocal feedback. **(H)** Example BLA rg.AAV injection site demonstrating ample TdTomato+ cells within, and neighboring, the BLA. This example mouse was selected despite AAV spread extending beyond the BLA to provide a strong test of any potential reciprocal innervation from the NAc or TuS to the BLA and surrounding amygdaloid nuclei. **(I)** Example image of the NAc and TuS in the mouse in (H) wherein no TdTomato+ cells are observed in either region supporting lack of feedback from either region onto the BLA. PCX = piriform cortex. Scale bars=500µm.

**Supplemental Figure 2. Co-expression of *drd1* and *drd2* is limited in the BLA and ventral striatum. Related to Figures 1 and 2. (A)** Example images of *drd1*+, *drd2*+, and an overlay of both *drd1*+ and *drd2*+ TuS neurons in a *drd1-RFP;drd2-GFP* BAC (bacterial artificial chromosome) double transgenic mouse, white arrowhead indicates a neuron that is *drd1+* and *drd2+*. Scale bars=50µm. **(B)** Quantification of *drd1*+, *drd2*+, or *drd1*+ and *drd2*+ neurons in the NAc, TuS, and BLA. n=3 mice, 3-5 sections/mouse/brain region. **(C)** Approach for defining whether, and if so, the extent of BLA🡪 ventral striatum contralateral spread wherein an AAV encoding synaptophysin.GFP in the BLA in one hemisphere and synaptophysin.mRuby in the BLA in the opposite hemisphere. **(D)** Representative images from one mouse of ipsilateral and contralateral projections from the BLA to the ventral striatum following injection as in (C). These example images show that while some BLA neurons transverse into the contralateral ventral striatum, qualitatively, the bulk of input is ipsilateral. Scale bars=500 µm. PCX= piriform cortex, NAcC= nucleus accumbens core, NAcSh= nucleus accumbens shell.

**Supplemental Figure 3. Synaptic properties of ventral striatum projecting BLA neurons. Related to Figure 3. (A)** and **(B)** are example current clamp recordings of *drd1+ and drd1*Ø ventral striatum neurons, respectively, showing current step driven action potentials. **(C)** and **(D)** are examples of light-evoked currents under voltage clamp recordings in *drd1+* and *drd1*Ø TuS neurons, in a *drd1-*Cre;Ai9 mouse. Light-evoked EPSCs were present after TTX and 4AP application, supporting action potential independent, ChR2-mediated monosynaptic transmission. EPSCs were blocked by bath application of 6-cyano-7-nitroquinoxaline-2,3-dione (CNQX) and d,l-2-amino-5-phosphonopentanoic acid (AP5), pointing towards glutamate as a principal transmitter which excites ventral striatum neurons. 4-AP = 4-aminopyridine, TTX = tetrodotoxin citrate, WO = wash out. The holding potential was -70 mV in C & D.

**Supplemental Figure 4. Mice display fear-related immobility and fear-related respiratory behavior during an odor-shock Pavlovian fear learning assay, reflective of associating odor with a mild shock only when the two are paired. Related to Figure 4. (Ai)** Schematic of three-day odor-shock Pavlovian aversion learning paradigm within a whole-body plethysmograph with both camera-captured immobility (from two cameras, 10 frames/sec) and pressure transducer captured respiratory behavior. The plethysmograph transducer allows for monitoring respiratory transients which were digitized along with stimulus delivery events. On day 1 the mice are habituated to handling and being in the plethysmograph. On day 2, the mice are delivered 10 trials of odor being paired or not with a mild aversive shock (0.5 mAmp foot shock via a metal floor, depicted by the lightning bolt). Finally, on day 3, the mice are delivered the odor again for 10 trials yet in the absence of shock for a test of memory/retrieval. **(Aii)** Individual trial schematic for paired and unpaired odor-shock Pavlovian fear learning, with the difference being that for the unpaired task the shock was delivered with a pseudorandom window during the inter-trial interval (ITI) so not to be associated with the conditioned odor. **(B)** Example data from a cohort of 56 c57bl/6j mice (roughly equal sexes in each condition). Increases in fear-related immobility from Trial 1 to Trial 10 during the learning day (Day 2) in mice receiving foot shock (left) (Two-way RM ANOVA *F*(4,51)=9.302, *p*<0.001), and sustained immobility in response to the odor paired with a foot shock on retrieval day (Day 3) (right) (One-way ANOVA *F*(4,51)=36.06, *p*<0.001). Significantly more time was spent in immobility by the paired group compared to unpaired, trial-shock, ITI-shock, or odor only groups. **(C)** Example respiratory traces corresponding to different behaviors, in which an upward deflection indicates an inhale and a downward deflection indicates an exhale. These examples are simply to help describe the variations in respiratory behavior, and how respiration is distinguished during stated of immobility (“freezing”). **(D)** Quantification of respiratory frequency in 8 c57bl/6j mice (roughly equal sexes in each condition), during the behaviors in (C). **(E)** Two-dimensional histograms of respiratory frequency aligned to odor delivery (black shaded box and dashed line) on the learning day (Day 2) during Trial 1 (top) and Trial 10 (bottom). **(F)** Quantification of the proportion of the odor delivery period (19s, excluding the last s wherein the shock occurred) spent in the 4-6Hz fear-related frequency range on the learning day (Day 2) during the odor presentation of Trial 1 vs Trial 10 (Two-way RM ANOVA *F*(4,49)=4.387, *p*=0.004). **(G)** Two-dimensional histograms of respiratory frequency aligned to odor delivery during retrieval (Day 3) on Trial 1 (top) and Trial 10 (bottom). **(H)** Quantification of the proportion of the odor delivery period (19s, excluding the last s wherein the shock occurred) spent in the 4-6Hz fear-related frequency range on retrieval day (Day 3) during all odor presentations (Trial 1-10) (One-way ANOVA *F*(4,49)=14.12, *p*<0.001). Mean±SEM.

**Supplemental Figure 5. TuS projecting *drd2*+ BLA neurons support aversion learning. Related to Figure 5.** **(A)** Influence of the DREADD agonist J60 (100nL, 10nM) on fear learning immobility in all NAc injected mice, **(C)** left NAc injected mCherry controls (*F*(1,27)=229, *p*<0.001); middle *drd1+* hM4D(Gi) mice (*F*(1,13)=155, *p*<0.001); and right *drd2+* hM4D(Gi) mice (*F*(1,13)=268, *p*<0.001). **(D)** Influence of J60 on immobility in all TuS injected mice, **(E)** left mCherry TuS injected controls (*F*(1,27)=252, *p*<0.001); middle *drd1+* hM4D(Gi) mice (*F*(1,13)=510, *p*<0.001); and right *drd2+* hM4D(Gi) mice (*F*(1,12)=133, *p*<0.001). All data analyzed using Two-way RM ANOVA, trial main effect reported. Mean±SEM.
